# SEED-MEDIATED RNA INTERFERENCE OF ANDROGEN SIGNALING AND SURVIVAL NETWORKS INDUCES CELL DEATH IN PROSTATE CANCER CELLS

**DOI:** 10.1101/2020.07.17.209379

**Authors:** Joshua M. Corbin, Constantin Georgescu, Jonathan D. Wren, Chao Xu, Adam S. Asch, Maria J. Ruiz-Echevarría

## Abstract

Resistance to anti-androgen therapy in prostate cancer (PCa) is often driven by genetic and epigenetic aberrations in the androgen receptor (AR) and coregulators that maintain androgen signaling activity. We show that specific small RNAs downregulate expression of multiple essential and AR-coregulatory genes, leading to potent androgen signaling inhibition and PCa cell death. Expression of different sh-/siRNAs designed to target TMEFF2, preferentially reduce viability of PCa, but not benign, cells and growth of murine xenografts. Surprisingly this effect is independent of TMEFF2 expression. Transcriptomic and sh/siRNA seed sequence studies indicate that expression of these toxic shRNAs lead to downregulation of AR-coregulatory and essential genes thru mRNA 3’-UTR sequence complementarity to the seed sequence of the toxic shRNAs. These findings reveal a specialized form of the Death Induced by Survival gene Elimination (DISE) mechanism in PCa cells, that we have termed Androgen Network (AN)-DISE, and suggest a novel therapeutic strategy for PCa.

## INTRODUCTION

Prostate cancer (PCa) is the most commonly diagnosed non-cutaneous malignancy and the second leading cause of cancer related deaths in men in the United States (Siegel, Miller, & Jemal, 2019). Androgen signaling is essential for normal prostate development and PCa cell growth and survival (Davey & Grossmann, 2016; Powers & Marker, 2013; Wang & Koul, 2017; Xie et al., 2017). This dependency is exploited in PCa treatment with androgen deprivation therapy (ADT) and targeted androgen receptor (AR) inhibitors (Davey & Grossmann, 2016). While these therapeutic modalities are initially beneficial, the majority of patients eventually relapse with a resistant and lethal form of the disease called castration-resistant PCa (CRPC) (Yuan et al., 2014). In most cases, CRPC cells retain AR expression and remain dependent on its activity for growth and survival, and adaptations, such as increased AR coregulator expression, AR amplification and/or mutation and constitutively active AR splice variants that lack the ligand binding domain, allow PCa cells to thrive in the androgen depleted environment (Huang, Jiang, Liang, & Jiang, 2018; Yuan et al., 2014). Second generation AR inhibitors and/or regimes that block adrenal androgen biosynthesis are often used to treat CRPC; however, the responses are short lived, leading to secondary resistance (Sharp, Welti, Blagg, & de Bono, 2016).

A novel mechanism, Death Induced by Survival gene Elimination (DISE), has recently been described as a potential cancer therapy (Putzbach, Gao, et al., 2018; Putzbach et al., 2017). DISE kills cancer cells through an RNA interference (RNAi) mechanism. In DISE, si- or shRNAs derived from CD95 and CD95L, or other human genes, function essentially as miRNAs to simultaneously target the 3’-UTR, and silence the expression of numerous essential genes, leading to cell death (Putzbach et al., 2017). DISE results in activation of multiple cell death pathways, which thwarts the development of resistance, and is preferentially toxic to transformed cells, including cancer stem cells (Ceppi et al., 2014; Hadji et al., 2014; Putzbach et al., 2017). Experiments with ovarian cancer orthotopic mouse models showed that DISE can be induced in vivo to kill cancer cells without evidence of off-site toxicity (Murmann et al., 2017).

In this study, we demonstrate that expression of certain shRNAs/siRNAs developed to target TMEFF2, an androgen regulated tumor suppressor gene with restricted expression to brain and prostate, promotes cancer cell death similar to DISE, independently of TMEFF2 targeting/expression. Distinct from DISE, in PCa cells, these shRNAs/siRNAs downregulate the AR and AR coregulatory genes, an effect that correlates with the presence of short seed matches in their 3’UTRs and which results in global androgen signaling inhibition and cell death. We have termed this mechanism Androgen Network DISE (AN-DISE). Both androgen dependent (AD) and CRPC cells are sensitive to AN-DISE, but different shRNAs/siRNAs have somewhat distinct effects on those cells, suggesting specificity. In vivo, AN-DISE shRNA significantly inhibit growth of CRPC 22Rv1 xenografts. These results suggest that AN-DISE may provide a potential therapeutic strategy for advanced PCa by simultaneously targeting the AR, multiple AR coregulators and, in essence, multiple essential pathways, making the development of resistance unlikely.

## RESULTS

### TMEFF2-targeted shRNAs reduce cell viability and growth PCa cells

We have previously reported a tumor suppressor function for TMEFF2 in PCa, with low TMEFF2 levels correlating with reduced disease-free survival (X. Chen et al., 2011; Corbin et al., 2016; Georgescu et al., 2019). To study the consequences of the loss of TMEFF2 expression in PCa, TMEFF2 was silenced in several PCa cell lines. Expression of five individual shRNAs to TMEFF2 (shTMEFF2-2, shTMEFF2-3, shTMEFF2-4, shTMEFF2-8 and shTMEFF2-9) spanning the length of the TMEFF2 transcript, decreased TMEFF2 expression (Figure 1 – figure supplement 1). Surprisingly expression of these shRNAs reduced growth and viability of PCa derived androgen dependent (LNCaP), castration resistant (22Rv1) and AR^-^ (DU145) cell lines when compared with the same cell lines expressing an shScramble control. shTMEFF2 also decreased growth and viability of other transformed cell lines, including pancreatic cancer Panc1 (not shown) and HEK293T-LX (Figure 1A and B). Importantly, these two cell lines and DU145 do not express TMEFF2, indicating that the effect on cell growth/viability is independent of shRNA-mediated silencing of TMEFF2.

**Figure 1.**
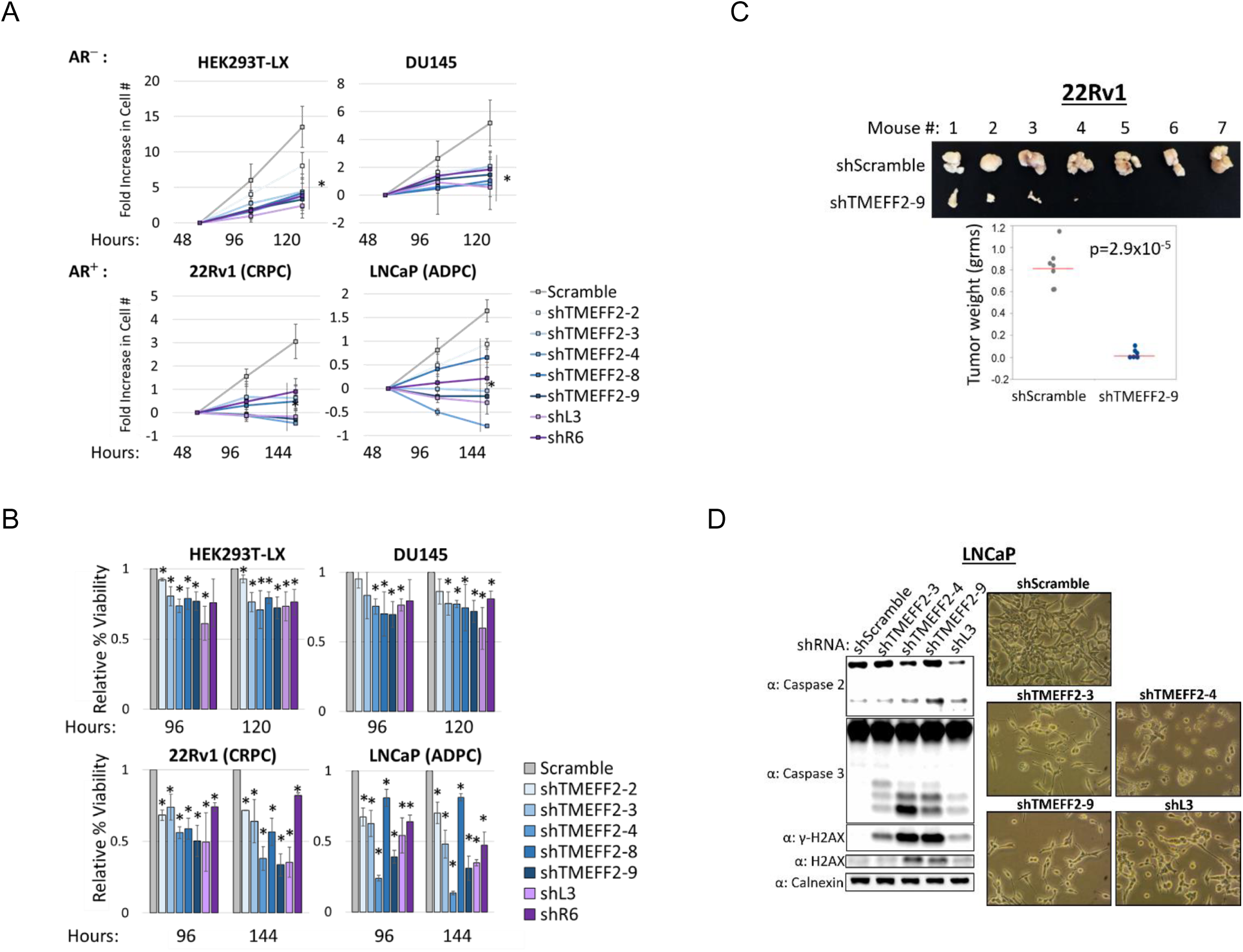
TMEFF2-targeted and DISE-inducing shRNAs reduce the growth and viability of transformed cells. (**A**) Cell growth and (**B**) viability of HEK293T-LX, DU145, 22Rv1 and LNCaP cells transduced with plasmids expressing TMEFF2-targeted shRNAs (shTMEFF2-2, -3, -4, -8, -9), CD95L and CD95 targeted shRNAs (shL3, shR6) or shScramble control. Cell growth is presented by fold increase in cell number relative to cell number at 48 hours after transductions. Viability was determined by trypan blue and is presented as percent viability relative to shScramble. N=4, error bars ±SD, * p<.05 determined by t-test. (**C**) Pictures of tumors (top panel) and dehydrated tumor weight (bottom panel) of xenograft tumors formed by 22Rv1 cells expressing dox-inducible shScramble and shTMEFF2-9 shRNAs after being injected subcutaneously into opposite flanks of NSG mice fed on doxycycline chow to induce shRNA expression. Mice were sacrificed and tumors were excised after four to five weeks of tumor growth. N=7, significance was determined by t-test for the means indicated by pink bar. (**D**) Western blot analysis (left panel) showing Caspase 2 cleavage, Caspase 3 cleavage, γ-H2AX and H2AX protein expression in lysates obtained from LNCaP cells expressing shTMEFF2-3, shTMEFF2-4, shTMEFF2-9, shL3 or shScramble control shRNA 96 hours after transduction. Pictures of cells (right panel) were taken 96 hours after transduction. Calnexin was used as a loading control.

In order to confirm that the effect on PCa cell viability is independent of TMEFF2 silencing, we used CRISPR Cas9 with two independent doxycycline (Dox)-inducible single guide RNAs (sgRNAs), to knockdown TMEFF2 expression in LNCaP cells. CRISPR-Cas9 knockdown of TMEFF2 (LNCaP-Cas9/sgTMEFF2) had no effect on cell viability as compared to the sgGFP control guide expressing cells (LNCaP-Cas9/sgGFP; Figure 1 – figure supplement 2A and B). However, expression of shTMEFF2-3, -4 or -9 in Dox induced LNCaP-Cas9/sgGFP or /sgTMEFF2 cells resulted in comparable loss in viability and increase in caspase 3 cleavage when compared to the corresponding cell lines expressing the shScramble control (Figure 1 – figure supplement 2C and D). This result confirmed that loss in PCa cell viability was not the result of TMEFF2 silencing, but of a mechanism dependent on expression of these TMEFF2 targeted shRNAs.

We then examined the effect of shRNA targeting TMEFF2 on the growth of subcutaneous tumors in mice. To this end, we established a Doxycycline responsive system using a lentiviral vector to express either shTMEFF2-9 or the shScramble in 22Rv1 (CRPC) cells. In cell culture, 5 days of Dox treatment was sufficient to knockdown TMEFF2 protein and induce Caspase 3 cleavage in shTMEFF2-9 expressing 22Rv1 cells compared to shScramble expressing cells (Figure 1 – figure supplement 3). For in vivo experiments, these two cell lines were grown in the absence of doxycycline –to prevent cell death, mixed with 50% basement membrane extract and inoculated subcutaneously into opposite flanks of NSG mice that were pre-fed (2 days) and kept in a doxycycline containing-diet for the duration of the experiment. Tumors were palpable/measurable in 100% of the mice 3 weeks after the injection with shScramble control cells but not in those mice injected with shTMEFF2-9 expressing cells. Mice were sacrificed ≈5 weeks after injections and tumors, if present, dissected and weighted. Expression of shTMEFF2-9 reduced or blocked tumor growth in this murine model (Figure 1C). Of note, the tumors that formed from shTMEFF2-9 expressing cells did not exhibit lower TMEFF2 levels compared to mouse-matched shScramble expressing tumors, as determined by IHC (not shown), suggesting that shTMEFF2-9 shRNA expression and/or activity may have been lost in these tumors.

The above results are reminiscent of DISE, a small RNA interference (RNAi)-mediated mechanism that preferentially kill cancer cells by targeting the 3′UTRs of multiple survival genes through a mechanism similar to miRNAs. In fact, known DISE-inducing shRNAs (shL3 targeting CD95L or shR6 targeting CD95 (Putzbach et al., 2017)) also reduced growth and viability of the five tested cell lines similar to the effect of shRNAs to TMEFF2 (Figure 1A and B). Moreover, similar to DISE, expression of shRNA to TMEFF2: i) induced DNA damage and apoptosis, as measured by histone H2AX phosphorylation and caspase 2/3 cleavage (Figure 1D); ii) promoted a phenotypic change that resulted in elongated, senescence-like enlarged cells (Figure 1D); and iii) prefentially affect viability of cancer cells, as observed by the attenuated effect on RWPE1, a normal prostate epithelial cell line (Figure 1 – figure supplement 4).

### TMEFF2-targeted shRNAs reduce androgen receptor expression and inhibit androgen response independently of TMEFF2 levels

As most PCa cells are critically dependent on AR signaling for growth, we hypothesized that these shRNA targeting TMEFF2 could also be targeting the AR and AR signaling in PCa cells. Using western blot analysis, we found that expression of nine independent shRNAs (shTMEFF2-1 – shTMEFF2-9) in LNCaP cells decreased TMEFF2 protein expression and resulted in reductions of AR and androgen responsive proteins PSA and FKBP5 particularly when grown in the presence of the AR ligand, dihydrotestosterone (DHT) (Figure 2A). Dose dependence experiments in LNCaP cells with two of those shRNAs (shTMEFF2-3 and shTMEFF2-9) demonstrated comparable reduction in TMEFF2 and PSA protein levels even at low concentrations of the shRNAs, with respect to the cells expressing the shScramble control (Figure 2 – figure supplement 1A and B). These results demonstrate that reduction in PSA level is not due to stress response or a consequence of high shRNA expression. Reductions in the level of androgen responsive proteins were also evident in C4-2B (CRPC) and 22Rv1 (CRPC) cells when TMEFF2-targeted shRNAs were expressed (Figure 2B, Figure 2 – figure supplement 2), suggesting that the inhibition of androgen signaling by TMEFF2-targeted shRNAs in PCa cells is not a cell line specific effect.

**Figure 2.**
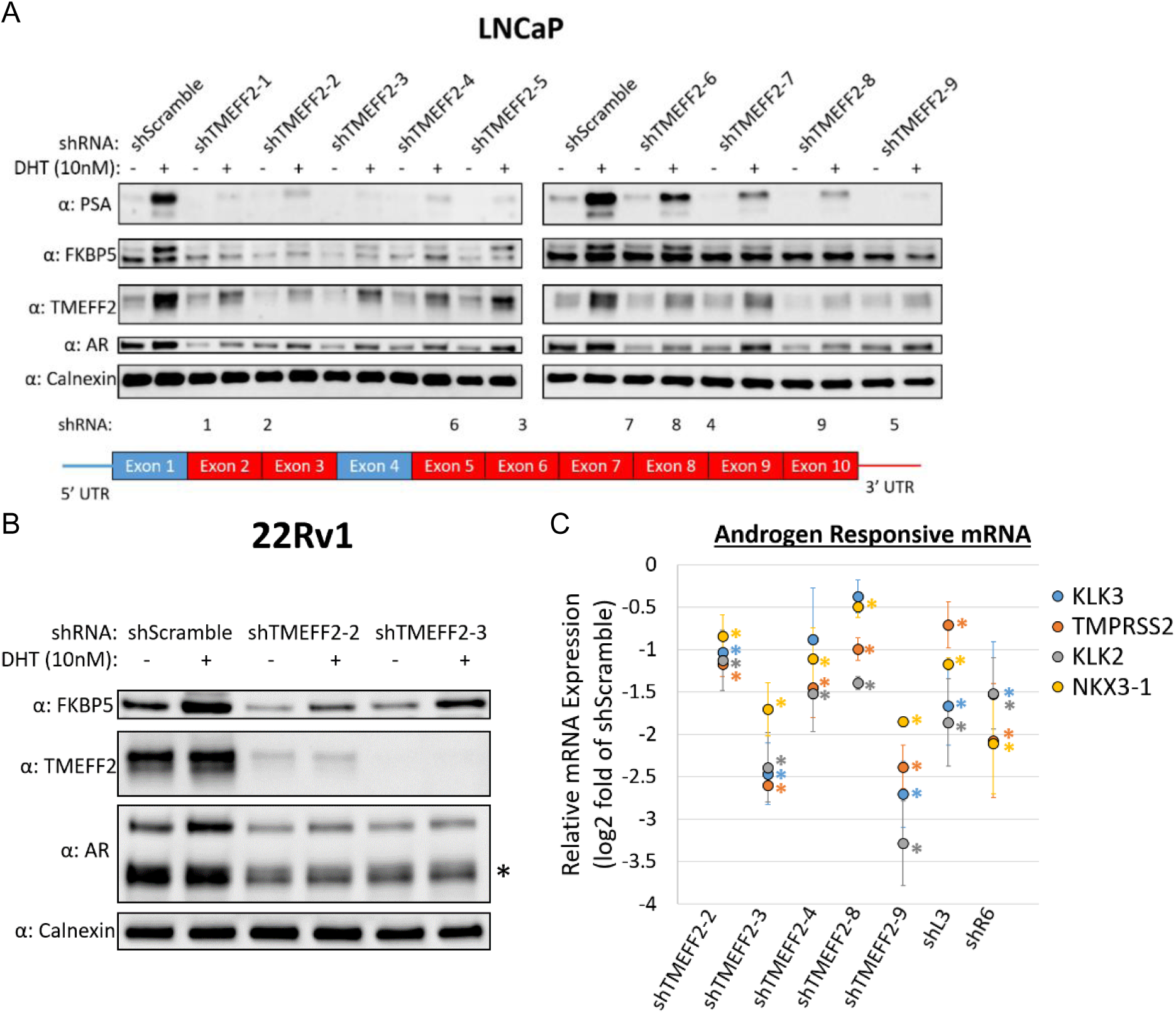
TMEFF2-targeted and DISE-inducing shRNAs inhibit androgen signaling in prostate cancer cells. (**A**) Western blot analysis showing protein levels of TMEFF2 (target gene), AR, and two androgen responsive proteins, PSA and FKBP5, in lysates from LNCaP cells expressing each of nine independent TMEFF2-targeted shRNAs or shScramble control, and grown in the presence and absence of 10 nM DHT. Calnexin was used for a loading control. The schematics in the lower panel indicate the exons targeted (red) or not targeted (blue) by the shRNAs, and the numbers corresponding to the shRNA names. (**B**) Western blot analyses showing TMEFF2, FKBP5 and AR (top band is full length AR, bottom band is the constitutively active AR-V7 isoform, see *) protein levels in 22Rv1 cells expressing shTMEFF2-2, shTMEFF2-3 or shScramble control, and grown in the presence and absence of 10 nM DHT. Calnexin was used for a loading control. (**C**) mRNA expression of androgen responsive genes, KLK3, TMPRSS2, KLK2 and NKX3-1 in LNCaP cells expressing TMEFF2-targeted shRNAs, shL3 or shR6 relative to cells expressing shScramble control. RNA was extracted 72 hours after transductions, and mRNA expression was determined by RT qPCR. RPL8, RPL38, PSMA1 and PPP2CA were the housekeeping genes used for normalization. N=3, error bars ±SD, * p<.05 determined by t-test.

We confirmed that the effect of shTMEFF2 RNAs on the levels of AR and androgen responsive proteins was independent of TMEFF2 levels by analyzing those proteins in lysates from Dox induced LNCaP-Cas9/sgGFP or /sgTMEFF2-1 cells that were subsequently transduced with plasmids expressing shTMEFF2-3 or shScramble. Western blot analysis indicated that AR, FKBP5 and PSA levels were similar in knockdown TMEFF2 cells (CAS9-sgTMEFF2-1) and in control cells (CAS9 sgGFP) expressing shScramble. Expression of shTMEFF2-3 drastically reduced AR, FKBP5 and PSA protein levels in both cell lines to a similar degree (Figure 2 – figure supplement 3A). Similarly, TMEFF2 knockdown in LNCaP cells using pooled antisense oligonucleotides (ASOs) did not alter PSA or AR protein levels despite reducing TMEFF2 levels as compared to cells transfected with the non-target control ASO (Figure 2 – figure supplement 3B). These data indicate that the effect of the TMEFF2-targeted shRNAs on androgen signaling was independent of their effect on TMEFF2 levels. In addition, deep RNA sequencing (RNA-seq) of the TMEFF2 locus indicated that no RNA originating from this locus could be a target of all nine TMEFF2 shRNAs, except for the main protein coding TMEFF2 isoform 1 mRNA (ENSEMBL ID ENST00000272771.9). Therefore, the TMEFF2 targeted shRNAs were not targeting a non-coding RNA that could be responsible for the phenotypes observed on androgen signaling and viability (Figure 2 – table supplement 1).

Finally, we also demonstrated that expression of shTMEFF2 in LNCaP cells reduced the mRNA expression of AR target genes (KLK3, KLK2, TMPRSS2, NKX3-1) to varying degrees, and in most cases the AR mRNA levels (Figure 2C, Figure 2 – figure supplement 4A). Interestingly, while DISE inducing shL3 also reduced the mRNA levels of some AR signaling targets, it did not affect AR mRNA or protein levels (Figure 2C, Figure 2 – figure supplement 4A and B), suggesting that androgen signaling inhibition was unlikely to be a sole consequence of reducing AR levels.

### TMEFF2 shRNAs reduce AR coregulatory and essential gene expression and inhibit global androgen response in LNCaP cells

The results above indicated that the effects on AR signaling and cell viability were completely independent of the presence or targeting of TMEFF2. To further understand the effect of shRNA to TMEFF2 on androgen signaling, we conducted RNA-seq with RNA from LNCaP cells transduced with shTMEFF2-3, -4, -9, or shScramble control shRNA and analyzed gene expression changes. In addition, we used the DISE inducing shL3. For these experiments, RNA was extracted 55 hours after transfection with the shRNAs, to circumvent potential changes in AR target levels secondary to loss of viability, which we observed only 72 hours after transfection (Figure Figure 3 – figure supplement 1). In fact, reductions in AR, KLK3 and TMPRSS2 mRNAs were evident at 48 hours, indicating that inhibition of androgen signaling is not a consequence of shRNA toxicity.

Genes that exhibited ± .5 log2 fold change and FDR q-value <.05 in samples from target shRNA expressing cells relative to shScramble control were considered to be significantly differentially expressed genes (DEGs). Pairwise comparisons of DEGs for the four shRNAs indicated a highly significant overlap (Figure 3 – figure supplement 2A and B), suggesting that the transcriptomic changes induced by the shRNAs were similar. Gene set enrichment analyses (GSEAs) (Mootha et al., 2003; Subramanian et al., 2005) identified 72, out of 19,695 tested MSigDB gene sets, that had significant differential expression (q-value < .25) in response to all four shRNAs (Figure 3A, Figure 3 – table supplement 1). All 72 of these common gene sets were downregulated by each of the shRNAs, and the most highly significant (q-value < .05) were Hallmark Androgen Response and Nelson Response to Androgen Up (Figure 3A, Figure 3 – table supplement 1). Importantly, 19 of the 100 Hallmark Androgen Response genes, including many prominent AR targets such as TMPRSS2, KLK2, NKX3-1 and FKBP5, were located in the leading edge of the GSEA enrichment plots for all four shRNAs, demonstrating a strong overlap among the most significantly downregulated androgen responsive genes (Figure 3A). All together, these results indicate that all four shRNA, including shL3, significantly affect global androgen signaling.

**Figure 3.**
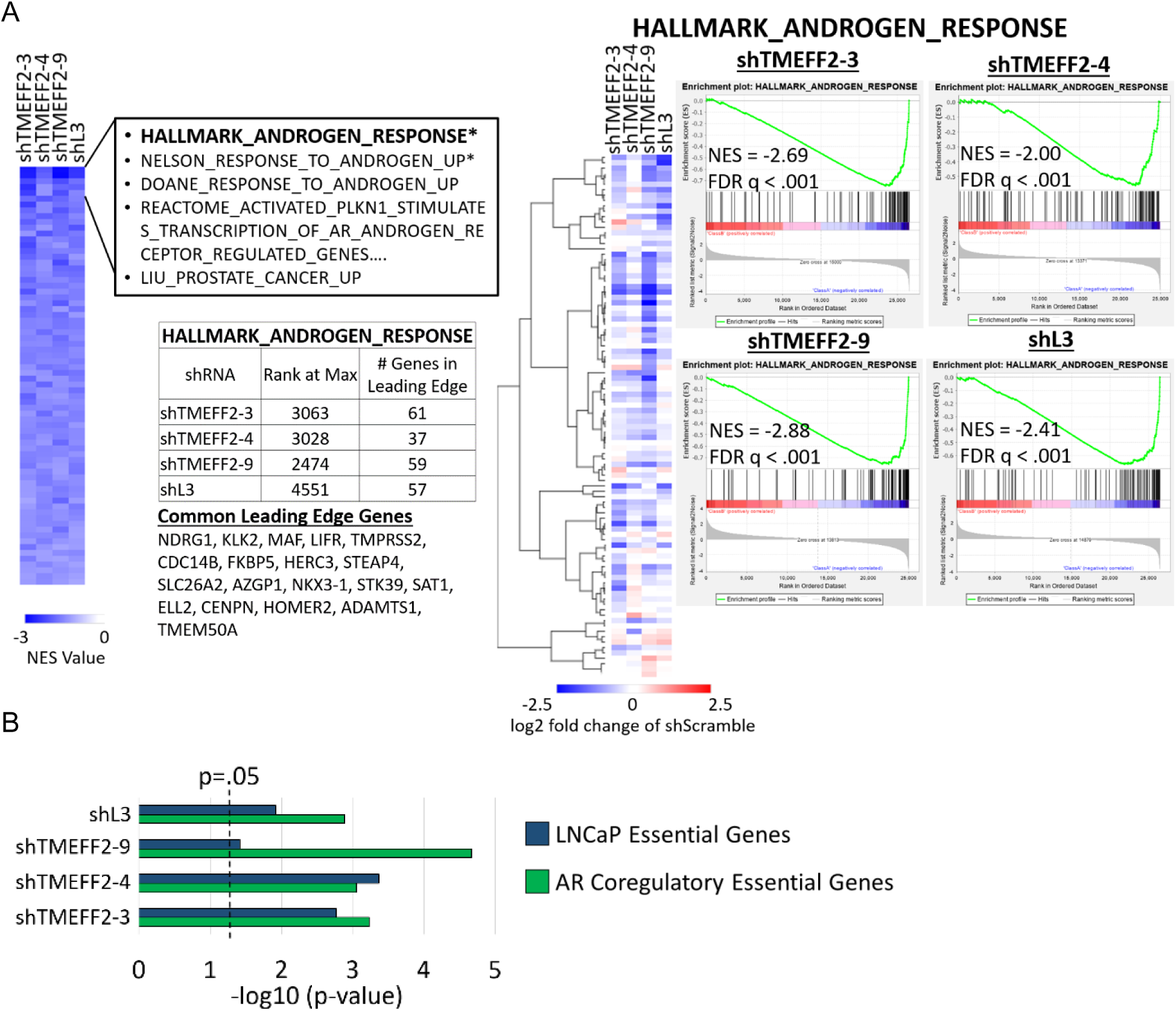
TMEFF2-targeted shRNAs and shL3 inhibit androgen transcriptional response and downregulate AR coregulatory and essential genes. (**A**) Heatmap (left panel) shows normalized enrichment scores (NES) of common significantly enriched gene sets (q-value <.25, for each shRNA) from gene set enrichment analyses (GSEA) of RNA-seq data from LNCaP cells expressing designated shRNAs compared to cells expressing the shScramble control. Negative NES indicates gene sets are enriched among downregulated genes, relative to shScramble control cells. Heatmap (middle panel) shows relative gene expression of genes within the Hallmark Androgen Response gene set. GSEA enrichment plots (right panel) show enrichment of the Hallmark Androgen Response gene set within gene lists rank ordered by gene expression (upregulated to downregulated) in cells expressing designated shRNAs relative to shScramble control cells. NES and p-value are labeled on each plot. The number of Hallmark Androgen Response genes within the leading edge for each shRNA are indicated in the table, and the 19 genes located in the leading edge for all shRNAs are listed. (**B**) –log10 p-value for enrichment of AR coregulatory genes (DePriest et al., 2016) and LNCaP essential genes (Fei et al., 2017) among significantly downregulated genes by each shRNA relative to shScramble expressing LNCaP cells. –log10 p-values are based on hypergeometric distribution.

Our results indicating that shTMEFF2 decrease the levels of AR mRNA and protein (Figure 2 – figure supplement 4A and B) could explain their effect on androgen signaling. However, DISE inducing shL3 affects androgen signaling without reducing AR levels. Critical to AR-mediated signaling is the recruitment of AR coregulators, which modulate its transcriptional response. We therefore examined the effect of the three TMEFF2 and the L3 targeted shRNAs on AR-coregulatory gene expression and determined that an AR-coregulatory gene set (n=274) (DePriest et al., 2016) was significantly enriched among genes downregulated by each of the 4 shRNAs (Figure 3B). Moreover, since cell viability was also affected in non-PCa cell lines (HEK293-LX) which are not dependent on AR signaling, we analyzed the effect of these 4 shRNAs on expression of essential genes. An essential gene set published for LNCaP cells (Fei et al., 2017) was significantly enriched among genes downregulated by each of the 4 shRNAs (Figure 3B). Finally, we observed that multiple histone genes were significantly downregulated by each shRNA (Figure 3 – figure supplement 3), consistent with previous reports of the transcriptomic effects of DISE (Putzbach et al., 2017).

To confirm the TMEFF2 shRNA-mediated influence on the androgen regulated transcriptome, RNA-seq was conducted with RNA extracted from shScramble and shTMEFF2-3 expressing LNCaP cells grown in the presence or absence of 10 nM dihydrotestosterone (DHT). RNA-seq analyses revealed that global androgen transcriptional response was nearly completely abolished in LNCaP cells by the expression of shTMEFF2-3 (13 DEGs in the presence vs. absence of DHT) compared to the shScramble (1,423 DEGs in the presence vs. absence of DHT) (Figure 3 – figure supplement 3A). GSEAs of MsigDB gene sets revealed that a large number of gene sets significantly modulated (up or down) by DHT in shScramble cells were regulated in the opposite direction by shTMEFF2-3 (Figure 3 - figure supplement 5, Figure 3 – table supplement 2 and 3). Strikingly, the normalized enrichment score (NES) values of these gene sets oppositely regulated by DHT and shTMEFF2-3 are significantly negatively correlated. The correlation was higher in the presence of DHT (R^2^= 0.6477 and R^2^= 0.1101 for and up- and downregulated sets) than in its absence (R^2^ = 0.2697 and R^2^ = 0.0045 for up- and downregulated sets) (Figure 3 – figure supplement 3A).

As expected, due to the androgen dependent nature of LNCaP cells, the essential gene set was significantly enriched among genes upregulated by DHT in shScramble expressing cells (Figure 3 – figure supplement 3B). However, essential and AR-coregulatory gene sets were significantly enriched among genes downregulated by shTMEFF2-3 in the presence and absence of DHT (Figure 3 – figure supplement 3B). Together, these data suggest that inhibition of androgen response is a major contribution to the TMEFF2 shRNA-mediated transcriptomic alterations. We have termed this mechanism in which DISE is triggered by global inhibition of androgen signaling, androgen-network (AN)-DISE.

### Downregulation of AR coregulatory and essential genes is associated with 3’UTR sequence complementarity to AN-DISE shRNA seed sequences

A key element of the DISE mechanism is RNAi seed mediated targeting of RISC to the 3’ UTR of essential genes, resulting in transcript downregulation. To begin understanding whether seed sequences are important in AN-DISE, we utilized the cWords software (Rasmussen, Jacobsen, & Krogh, 2013) to identify sequences enriched in the 3’-UTRs of downregulated mRNAs (from RNA-Seq data) in cells transfected with shTMEFF2-3, -4, -9, or shL3, and then determined whether the enriched sequences were complementary to the seed sequence from the mature guide strand of the different shRNAs. cWords assesses over-representation of nucleotide words in fold-change expression ranked ordered gene lists, correlating differential expression and motif occurrence (Rasmussen et al., 2013). For each of the shRNAs, the most significantly enriched 6-8 nucleotide sequences in the 3’ UTR of downregulated genes were complementary to potential seeds and surrounding nucleotides on the guide strand (Figure 4A, Figure 4 – figure supplement 1, Figure 4 – figure supplement 2, Figure 4 – figure supplement 3, Figure 4 – figure supplement 4 and Figure 4 – figure supplement 5). Sequences within coding regions complementary to shRNA seeds exhibited a weak association with gene downregulation (Figure 4 – figure supplement 2, Figure 4 – figure supplement 3 and Figure 4 – figure supplement 5). Furthermore, in each case, the most significantly enriched 6mer sequence was complementary to the seed sequence of the shRNA guide strand that would be produced with a Dicer cut after nucleotide +3 during shRNA processing. This corresponds with a highly predicted Dicer cut site (Putzbach et al., 2017). I*n silico* data from a previously published seed sequence toxicity screen (Gao et al., 2018), indicates the toxicity of these seeds. In fact, using this data, 7 out of 9 TMEFF2 targeted shRNAs were found to contain 6mer seeds within the 50^th^ percentile for reduced viability out of all 4096 possible 6mer seeds (Figure 4 – figure supplement 6).

**Figure 4.**
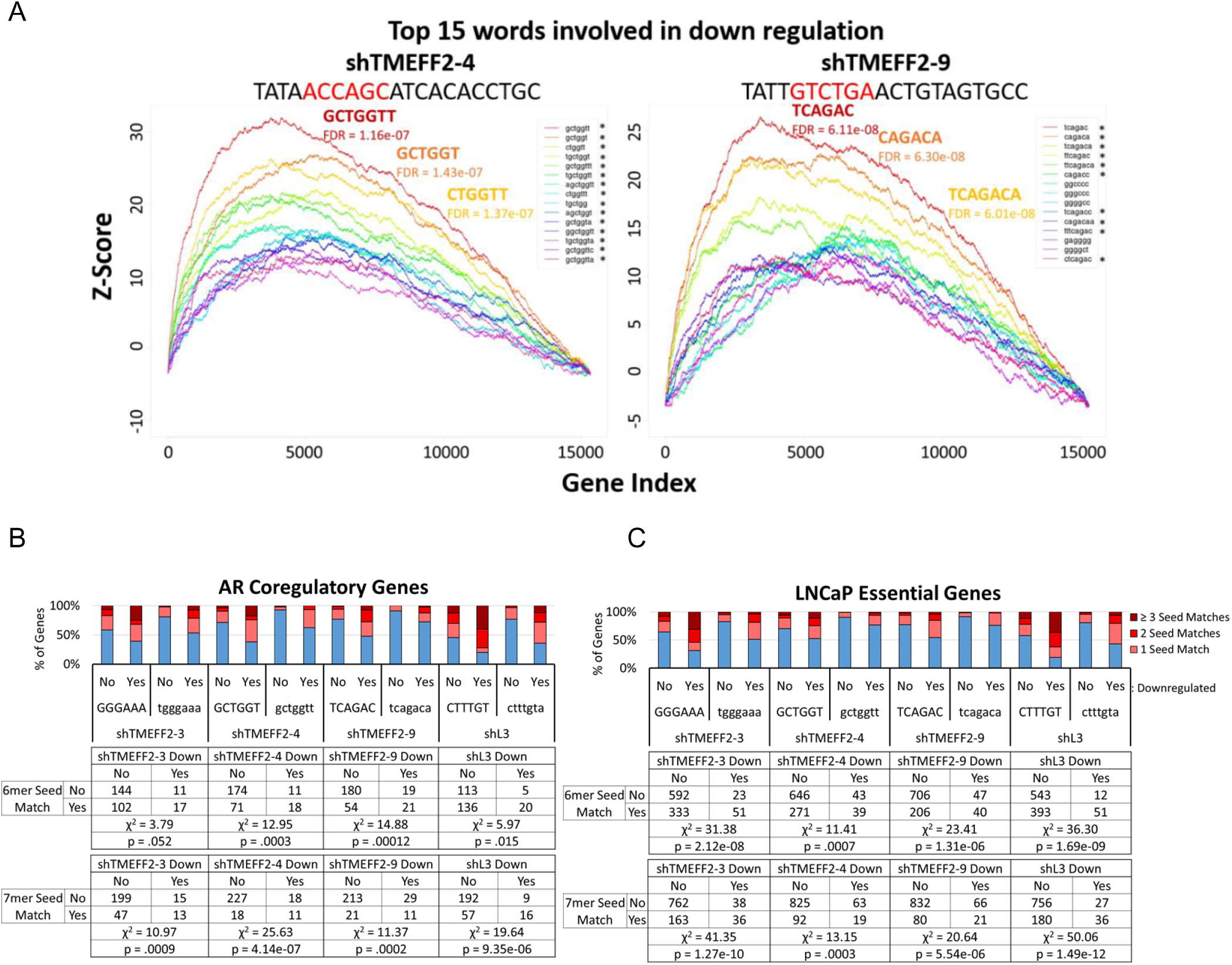
AR coregulatory and essential gene downregulation are associated with 6mer and 7mer 3’ UTR sequences complementary to potential shRNA seed sequences. (**A**) cWords enrichment plots showing the top 15 enriched 6mer, 7mer and 8mer in the 3’ UTR of genes downregulated by designated shRNAs according to RNA-seq analyses. Y-axis shows Z-score enrichment values. X-axis contains rank ordered genes from the most downregulated to upregulated expression. The top 3 enriched 3’ UTR sequences and FDR values are labeled (red: most enriched, orange: second most enriched, yellow: third most enriched). Unprocessed shRNA guide strand sequences are above each plot, and the potential 6mer seed sequence complementary to the most enriched 6mer 3’ UTR sequence associated with gene downregulation is in red. * indicates enriched sequences that are complementary to potential shRNA seed motifs. (**B**) AR coregulatory (DePriest et al., 2016) and (**C**) LNCaP essential gene (Fei et al., 2017) sets stratified by downregulation (yes or no) and by the presence in their 3’-UTR of single, 2, or more, 6mer (upper case) or 7mer (lower case) sequences identified by cWords analyses. Only the most significantly associated with gene downregulation by each shRNA (potential 3’ UTR seed matches) were used. Contingency tables are located below each stacked bar graph. P-values were calculated by chi square test of independence, and are labeled on the bottom of each contingency table.

Our analysis with cWords also identified endogenous miRNAs that share similar seed motifs with each shRNA, and miRNA Database (miRDB, mirdb.org) predicted target gene sets for these miRNAs were also significantly downregulated by the TMEFF2 or L3 shRNAs containing the corresponding similar seed (Figure 4 – figure supplement 7). Interestingly, the seed sequence identified for shTMEFF2-4 is identical to the seed sequence of miR-634, which has been shown to induce apoptosis in cell lines of multiple cancer types (Fujiwara et al., 2015; Gokita, Inoue, Ishihara, Kojima, & Inazawa, 2020; Zhang, Cao, Fu, Yun, & Zhang, 2016), including PCa cell lines, and to reduce AR protein levels and viability in PCa cells (Ostling et al., 2011). These results suggest an association between shRNA seed complementarity and gene expression downregulation.

We next tested the association between shRNA seed complementarity and downregulation of known AR coregulatory (DePriest et al., 2016) and essential genes (Fei et al., 2017). Using RNA-seq data for each of the shRNAs (shTMEFF2-3, -4, -9 and L3), we determined the correlation between the absence or presence of single or multiple sequences complementary to the seed sequence of each shRNA in the 3’-UTR of AR-coregulatory and essential genes, and their downregulation in LNCaP cells expressing the corresponding shRNAs. For all four shRNAs, 6mer and 7mer seed complementarity in the 3’ UTR was significantly associated with AR coregulatory and essential gene downregulation (Figure 4B and C). RT qPCR experiments were also used to validate some of these results (Figure 4 – figure supplement 8). A similar analysis was conducted for the list of essential genes using RNA-seq data from LNCaP cells expressing shTMEFF2-3 and shScramble grown in the presence and absence of DHT. Since a significant number of androgen induced essential genes were downregulated by shTMEFF2-3, we stratified the essential genes based on whether they are androgen induced (n=66) or not (n=933). In this case, the association with 3’-UTR complementarity was significant only for the non-androgen induced group (Figure 4 – figure supplement 9). These results suggest that a large number of androgen induced essential genes were downregulated indirectly by shTMEFF2-3 mediated androgen signaling inhibition. Based on our previous observations, this likely results from seed-mediated AR coregulatory gene downregulation (of note, 21 out of the 274 known coregulators have 3’-UTR matches to the shTMEFF2-3 6-mer seed and are downregulated in the presence and/or absence of DHT).

### AN-DISE mediated effects on AR coregulatory gene downregulation, androgen signaling inhibition and PCa cell viability are seed mediated

To further define the relevance of the seed sequence in the AN-DISE phenotype, we used AN-DISE siRNAs (siTMEFF2-3, siTMEFF2-4, siTMEFF2-9) that had matching mature guide strands, and therefore seed sequences, to shTMEFF2-3, shTMEFF2-4 and shTMEFF2-9, and measured their effect on androgen signaling and cell viability when transfected in LNCaP cells. In addition, we designed several controls: i) the same siRNAs carrying 5’ ON-TARGET-Plus modifications (Dharmacon, GE Healthcare), which block seed-mediated off-target effects (5p-siTMEFF2). ii) siTMEFF2-4+3 and siTMEFF2-4+5 siRNAs that have a shifted siTMEFF2-4 target sequence by 3 nucleotides and 5 nucleotides respectively, and therefore, while still fully complementary to TMEFF2, they have a different seed sequence than siTMEFF2-4, and iii) a miR-634 siRNA mimic (si634) which contained the same seed sequence as siTMEFF2-4 independent of the full TMEFF2 target sequence. All three TMEFF2-targeted siRNAs (siTMEFF2-3, -4, -9) significantly reduced LNCaP viability when compared to the siNon-target control, while transfection of the ON-TARGET-Plus modified TMEFF2-targeted siRNAs had a much reduced effect on viability (Figure 5A, Figure 5 – figure supplement 1). Shifting the siTMEFF2-4 target sequence (siTMEFF2-4+3 and shTMEFF2-4+5) had a much weaker effect on viability. Importantly, si634 exerted similar toxicity to its non-modified “seed homolog” siTMEFF2-4 without significantly reducing TMEFF2 expression (Figure 5A). The effect on viability of the different siRNAs correlated with Caspase 3 cleavage, but not with their ability to silence TMEFF2 expression (Figure 5A and B, Figure 5 – figure supplement 1). These results indicate that AN-DISE toxicity in PCa cells is dependent on the seed mediated targeting mechanism of the siRNAs, leading to apoptosis. These experiments also confirm that the observed toxicity is not due to an interferon-like mechanism or TMEFF2 silencing. RT-qPCR analysis of the AR, AR-responsive and AR-coregulatory genes, in cells transfected with each of the siRNAs listed above showed that those siRNAs that more greatly affect viability, also more potently downregulate expression of those targets. These data indicate that AR coregulatory gene downregulation, androgen signaling inhibition and loss in PCa cell viability are correlated and RNAi seed-mediated (Figure 5C, Figure 5 – figure supplement 2).

**Figure 5.**
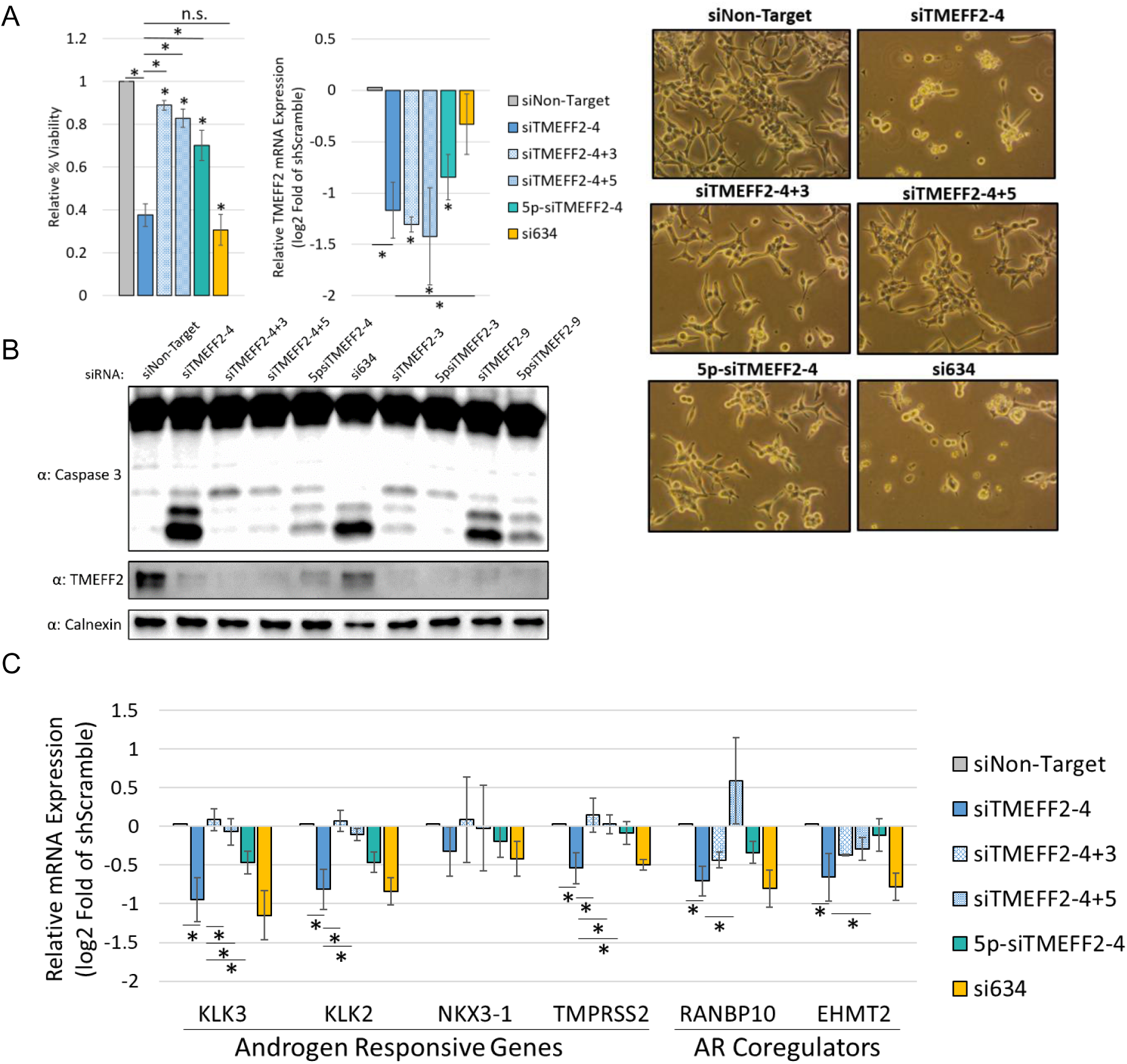
AR coregulatory gene downregulation and decrease in PCa cancer cell viability are shRNA seed mediated. (**A**) Relative percent viability and TMEFF2 mRNA expression in LNCaP cells transfected with siTMEFF2-4 and control siRNAs (siNon-Target; Negative seed controls: siTMEFF2-4+3, siTMEFF2-4+5, 5p-siTMEFF2-4; Positive seed control: si634). 5p designates an ON-Target-Plus modification (Dharmacon) that blocks seed mediated gene downregulation. Cell viability was determined by trypan blue. mRNA expression was determined by RT qPCR using RPL8, RPL38, PSMA1 and PPP2CA housekeeping genes for normalization. Viability measurements, cell pictures and RNA extractions were done 72 hours after siRNA transfections N=4, error bars ±SD, * p<.05 compared to siNon-target. Bars with * designate significant differences relative to siTMEFF2-4. Significance was determined by t-test. (**B**) Western blot analysis showing Caspase 3 cleavage and TMEFF2 protein expression in lysates from LNCaP cells transfected with the designated siRNAs. Lysates were obtained 72 hours after siRNA transfections. Calnexin was used as a loading control. (**C**) Relative androgen responsive, AR coregulator and AR mRNA expression in LNCaP cells transfected with siTMEFF2-4 and control siRNAs. RNA was extracted 72 hours after siRNA transfections. mRNA expression was determined by RT qPCR using RPL8, RPL38, PSMA1 and PPP2CA housekeeping genes for normalization. N=4, error bars ±SD, * p<.05 determined by t-test.

In agreement with previous reports that DISE is a mechanism that is preferentially active in cancer cells, transfection of Cy5 labeled siTMEFF2-4 and siTMEFF2-9 in normal prostate epithelial cell lines did not affect viability when compared to the Non-target control. However, these siRNAs significantly reduce viability of the LNCaP, C4-2B and 22Rv1 PCa cell lines (Figure 5 – figure supplement 3), consistent with reports that DISE preferentially occurs in cancer cells (Hadji et al., 2014).

In summary, the data presented in this study suggest that certain RNAi seed sequences can induce AN-DISE, a form of DISE that is specific to PCa cells. In AN-DISE essential and AR coregulatory genes are downregulated through an RNAi seed-mediated mechanism, leading to potent androgen signaling inhibition, downregulation of androgen regulated survival pathways and PCa cell death (Figure 6).

**Figure 6.**
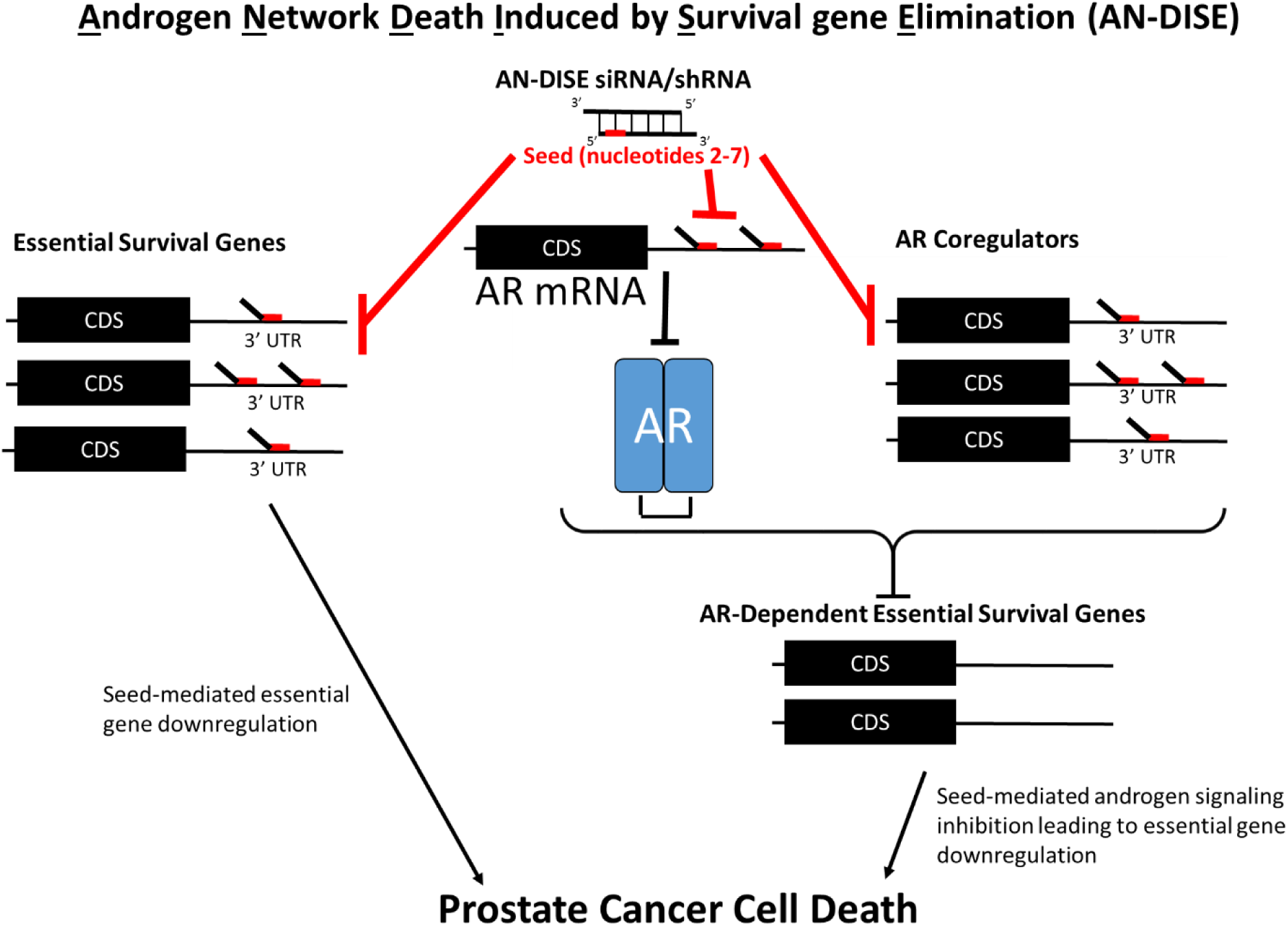
AN-DISE model. Seed sequences of certain RNAi molecules in PCa cells lead to : i) seed-mediated downregulation of essential genes and ii) androgen signaling regulatory genes leading to androgen signaling inhibition and downregulation of androgen regulated survival pathways.

## DISCUSSION

Therapeutic resistance to ADT in PCa remains a major hurdle in treatment (Huang et al., 2018; Yuan et al., 2014). Various molecular mechanisms underlie the persistent activity of the AR in CRPC cells growing in an androgen depleted environment, including AR mutations and splice variants, and the overexpression of AR coregulators (Yuan et al., 2014). Furthermore, the AR cistrome and transcriptional program undergoes changes that increase oncogenicity during the progression to CRPC (Z. Chen et al., 2018; Decker et al., 2012), and multiple AR coregulators may play significant roles in this process (Z. Chen et al., 2018; Groner et al., 2016; Xu et al., 2012). However, identifying individual essential coregulators that can be effectively targeted in the clinical setting has proved difficult.

In this study, we show that RNAi can be used in PCa cells to target and downregulate multiple AR coregulatory and essential gene networks, through a miRNA-like mechanism, resulting in androgen signaling inhibition and PCa cell death. Since androgen signaling inhibition via the seed mediated downregulation of AR coregulators plays a central role in the cell death mechanism, and RNAi-induced transcriptomic changes show profound effects on androgen regulated genes, we term this RNAi mechanism, which resembles DISE (Putzbach, Gao, et al., 2018; Putzbach et al., 2017), androgen network DISE (AN-DISE). We independently demonstrate that in addition to the novel shRNAs identified in this study, certain shRNAs previously shown to induce DISE in other cancer cell types induce AN-DISE in PCa cells. Interestingly, some AN-DISE RNAi molecules promote DISE-like cell death in non-AR signaling dependent cells. This is supported by the observation that AN-DISE induced shRNAs reduce growth/viability of AR-negative DU145 and HEK293T-LX cells. The centrality of androgen regulated survival genes in PCa and their relationship to survival gene pathways in cells lacking the androgen receptor raise the possibility of an evolutionary relationship between the development of cell survival networks and hormonal signaling. Our data also clearly show that different shRNAs display a certain level of specificity with respect to the targeting of distinct AR-coregulatory genes.

The interpretation of loss of function experiments using RNAi is often clouded by the miRNA-like seed mediated targeting of RISC to the 3’ UTRs of multiple genes, resulting in transcript downregulation and off-target effects (Jackson et al., 2006; Jackson & Linsley, 2010; Kamola, Nakano, Takahashi, Wilson, & Ui-Tei, 2015). In fact, considering potential off-target effects has aided in the interpretation of RNAi screen studies (Buehler et al., 2012; Riba et al., 2017; Sudbery, Enright, Fraser, & Dunham, 2010). Clinically, the majority of RNAi-based cancer therapies that have entered clinical trials have been designed to target individual genes (Davis et al., 2010; Schultheis et al., 2014; Tabernero et al., 2013). However, strategies based on the seed-mediated targeting of essential gene networks provide many theoretical advantages, including a reduction in evolutionary avenues for adaptive resistance in cancer cells. This is supported by a recently described DISE mechanism (Putzbach, Gao, et al., 2018; Putzbach et al., 2017). DISE works through the RNAi seed-mediated downregulation of numerous essential genes leading to cancer cell death through activation of multiple death pathways (Hadji et al., 2014; Putzbach et al., 2017) making adaptive resistance much less likely when compared to strategies targeting individual genes. Importantly, preclinical studies have shown that DISE-inducing siRNAs can be delivered to ovarian tumors in mice via nanoparticles, and can significantly reduce tumor growth (Murmann et al., 2017).

We propose that AN-DISE represents a potential therapeutic strategy that has important theoretical benefits over conventional androgen signaling targeted PCa therapies:

1. Ability to downregulate multiple AR coregulators. The simultaneous downregulation of multiple AR coregulators with a single therapeutic agent bypasses the necessity for identifying single targetable essential coregulators, which may vary according PCa subtype, tumor cell heterogeneity, and may change during PCa progression. In addition, the consequences of functional redundancies of AR coregulators are likely overcome by targeting multiple AR coregulators simultaneously.
2. Ability to target constitutively active AR isoforms. AN-DISE induces CRPC cell death, and certain seed sequences downregulate AR mRNA and/or protein, including AR-V7, which is associated with increased resistance to ADT and AR inhibitors (Antonarakis et al., 2014).

Of note, since AN-DISE RNAi also downregulates multiple essential genes that are independent of the AR signaling axis, the risk of developing AR-independent resistance mechanisms, common to treatment with second generation anti-androgens, is likely reduced.

The presence of DISE-inducing RNAi targeting in certain genes, such as CD95 and CD95L, (Patel & Peter, 2018; Putzbach et al., 2017) and TMEFF2 (this study), that can have a tumor suppressor role, may point to an endogenous tumor suppressive mechanism in which small RNAs can be processed from certain mRNAs and loaded into RISC, resulting in essential gene downregulation. In support of this hypothesis, a previous report suggests that small RNAs derived from CD95L mRNA can be loaded into RISC and induce an endogenous DISE mechanism in cancer cells. Interestingly, secondary structure predictions indicate that CD95L mRNA forms a tightly folded structure with extensive regions of complementarity that could provide the endogenous double stranded sequences to be processed and loaded into RISC (Putzbach, Haluck-Kangas, et al., 2018). A similar prediction was obtained for the TMEFF2 mRNA (data not shown). This raises questions of whether DISE can be induced endogenously in cancer cells, and whether the endogenous mechanism could be exploited therapeutically without the exogenous delivery of RNAi. Furthermore, if TMEFF2 mRNA can function as tumor suppressor through this mechanism, then it is possible that both TMEFF2 protein and mRNA function independently as tumor suppressors. We have previously published that low TMEFF2 mRNA expression correlates with decreased disease-free survival in PCa patients (Georgescu et al., 2019). Since TMEFF2 expression is induced by the AR (Gery, Sawyers, Agus, Said, & Koeffler, 2002; Overcash et al., 2013), TMEFF2 mRNA may be part of a negative feedback loop with androgen signaling through AR coregulator downregulation via small TMEFF2-derived RNAs being loaded into RISC. This mechanism could exert pressure on PCa cells to lose TMEFF2 expression during PCa progression. The existence of endogenous DISE and AN-DISE mechanisms warrant further investigation.

In summary, in addition to supporting previous reports describing the DISE mechanism, we describe a novel form of DISE in PCa cells, AN-DISE. By downregulating multiple essential genes and AR coregulatory genes through an RNAi seed-based mechanism, we propose that AN-DISE represents a potential therapeutic strategy for inhibiting androgen signaling and inducing PCa cell death.

## MATERIALS AND METHODS

### Cell culture and plasmid constructs

LNCaP, 22Rv1, DU145 and RWPE1 (CRL11609™) cell lines were obtained from American Type Culture Collection (ATCC^®^, Manassas, VA) and maintained at low passage. BHPre1 and NHPre1 cells were obtained from Dr. S. Hayward (NorthShore Research Institute), and LentiX-293T cells were obtained from Clontech/Takara. LNCaP and 22Rv1 cells were maintained in RPMI Glutamax growth media (Gibco). DU145, PC3 and LentiX-293T cells were maintained in DMEM growth media (Gibco). Both RPMI and DMEM media were supplemented with 10% fetal bovine serum (FBS) (Corning), 100 units/mL penicillin, 100 μg/mL streptomycin, Amphotericin B and 2mM L-Glutamine. RWPE1 cells were maintained in KSF media (Gibco). BHPre1 and NHPre1 cells were maintained in HPrE-conditional medium, as previously described (Jiang et al., 2010). Cells were tested negative for mycoplasma using the Mycosensor PCR assay kit (Agilent) or the LookOut Mycoplasma PCR detection kit (Sigma-Aldrich).

plasmid pLKO.1 vector was used for shRNA expression, and plasmids were obtained from Open Biosystems or cloned using pLKO.1-TRC (a gift from Dr. David Root (Addgene # 10878; http://n2t.net/addgene:10878; RRID: Addgene 10878)). For doxycycline inducible expression, shRNAs were cloned into Tet-pLKO-Puro (a gift from Dmitri Wiederschain (Addgene plasmid # 21915; http://n2t.net/addgene:21915; RRID: Addgene_21915)). pLKO.1-TRC cloning protocol was obtained from addgene. shTMEFF2-6 and shTMEFF2-9 sequences were generated using the siRNA Wizard online tool by Invitrogen: https://www.invivogen.com/sirnawizard/). All other shRNAs used in this study come from the RNAi consortium shRNA library, and TRC numbers are provided.

shScramble (CCTAAGGTTAAGTCGCCCTCG)
shTMEFF2-1 (CTGGTTATGATGACAGAGAAA) TRCN0000073522
shTMEFF2-2 (CGTCTGTCAGTTCAAGTGCAA) TRCN0000222559
shTMEFF2-3 (GCGCTTCTGATGGGAAATCTT) TRCN0000073521
shTMEFF2-4 (GCAGGTGTGATGCTGGTTATA) TRCN0000073520
shTMEFF2-5 (CCTTGCATTTGTGGTAATCTA) TRCN0000073518
shTMEFF2-6 (GGCTCTGGAGAAACTAGTCAA)
shTMEFF2-7 (ATGCAGAGAATGCTAACAAAT) TRCN0000373776
shTMEFF2-8 (CATACCTTGTCCGGAACATTA) TRCN0000373700
shTMEFF2-9 (GGCACTACAGTTCAGACAATA)
shL3 (ACTGGGCTGTACTTTGTATAT) TRCN0000059000
shR6 (CCTGAAACAGTGGCAATAAAT) TRCN0000038696

CRISPR Cas9 (pRCCH-CMV-Cas9-2A-Hygro) and doxycycline inducible sgRNA (pRSGT16-U6Tet-(sg)-CMV-TetRep-2A-TagRFP-2A-Puro) plasmids were obtained from Cellecta. LentiX 293T cells and CalPhos transfection reagents (Clonetech) were used for viral particle packaging. psPAX2 and VSV-G plasmids were used for lentiviral packaging. psPAX2 was a gift from Dr. Didier Trono (Addgene plasmid # 12260; http://n2t.net/addgene:12260; RRID: Addgene_12260), and pCMV-VSV-G was a gift from Dr. Robert Weinberg (Addgene plasmid # 8454; http://n2t.net/addgene:8454; RRID: Addgene_8454). Viral concentrations necessary for approximately 90% or greater survival (for non-toxic constructs) in selection antibiotic were used for transductions. For lentiviral transductions, cells were seeded in 6 cm plates at 50% confluency, and viral particle containing supernatant was diluted in 1.5 ml 8ug/ml polybrene serum free media and added to cells. After 5 hours, 1.5 ml of 10% FBS growth media was added to transduction media. Growth media was refreshed 24 hours after initial viral particle exposure. When cell lines were stably selected, transduced cells were grown for an average of 10 days using the following selection antibiotic concentrations: Puromycin 1ug/ml, hygromycin 750ug/ml.

### Cell growth and viability assays

For growth and viability analyses in response to shRNA expression, LNCaP, 22Rv1, DU145 and HEK293T-LX (LentiX-293T, HEK293T subclone) cell lines were transduced with pLKO.1 shRNA constructs. Cells were trypsinized and seeded in 6 well plates 24 hours post-transduction at the following concentrations: 1×105 cells/well (HEK293T-LX, DU145), 2×105 cells/well (LNCaP, 22Rv1). Trypan blue was used to stain the cells to selectively count live cells and assess viability using a Nexcelom Auto T4 Cellometer. Cells were trypsinized and counted using the same method 24, 48, 72 (for DU145, HEK293T-LX) or 96 (for LNCaP, 22Rv1) hours after seeding the for initial cell count (48h). Of note, LNCaP and 22Rv1 have a slower growth rate. Viability was assessed at each time point via trypan blue. Viability of RWPE1 cells transduced with shRNA was determined 120 hours after transductions. Relative percent viability was then calculated by dividing percent viability of knockdown cell lines by percent viability in cells expressing the shScramble control.

For viability analyses in response to siRNA, transfections were carried out as described in the siRNA transfections section of the materials and methods. Cells were trypsinized and split 24 hours after transfections, and viability was determined by trypan blue 72 hours after transfections.

Statistics: Two-tailed T tests were used to calculate significant differences in viability and growth. P<.05 was considered to be statistically significant.

### Antisense oligo transfections

FANA antisense oligos (ASOs) were obtained from AUM Biotech. LNCaP cells were transfected with 250 nM ASO (Non-target or pool of 4 targeting TMEFF2) using .19% Dharmafect #3 transfection reagent. 48 hours after transfection, cells were treated with 10% CSS RPMI for 24 hours, followed by 24 hours in 10 nM dihydrotestosterone (DHT) or .0001% ethanol (EtOH).

### siRNA transfections

Custom siRNAs with and without the ON-TARGET-Plus modification to block seed mediated off-target effects were obtained through Dharmacon (GE Healthcare). When indicated custom siRNAs were also ordered with a Cy5 label on the 3’ end of the passenger strand. siRNA guide strand sequences were as follows:

siTMEFF2-3: AUUUCCCAUCAGAAGCGCAUU
siTMEFF2-4: AACCAGCAUCACACCUGCAUU
siTMEFF2-4+3: UAUAACCAGCAUCACACCUUU
siTMEFF2-4+5: AGUAUAACCAGCAUCACACUU
siTMEFF2-9: UGUCUGAACUGUAGUGCCCUU
si634: AACCAGCACCCCAACUUUGUU

Non-target siRNA pool (Dharmacon D-001810-10-05) and Cy5 labelled Non-target siRNA (same sequence as D-001810-01-05) were used as negative controls.

Dharmafect siRNA transfection protocol was used for transfections, (https://horizondiscovery.com/-/media/Files/Horizon/resources/Protocols/basic-dharmafect-protocol.pdf).Dharmafect reagent #1 (0.2%) was used for RWPE1, BHPre1 and NHPre1 cell lines, and 0.2% Dharmafect reagent #3 was used for LNCaP, C4-2B and 22Rv1 cell lines. 30 nM siRNA was used for each transfection.

### RNA extractions and lysate preparations

RNeasy (Qiagen) with on-column DNAse treatment (Qiagen) was used for RNA extractions. Cell Signaling lysis buffer 9803 was used for whole cell lysates. Cells were maintained in complete media containing 10% FBS for all experiments not using DHT. For experiments measuring gene expression response to DHT, cells grown after transduction with the shRNA were washed with PBS and treated with 10% charcoal stripped serum (CSS) containing growth media for 24 hours for hormone depletion, followed by 24 hours in the same media containing 10 nM DHT (Sigma Aldrich) or .0001% EtOH vehicle.

### Western blot analysis and antibodies

Proteins were separated via 10% SDS/PAGE using mini-PROTEAN TGX stain free gels (Biorad), and transferred onto .2 um nitrocellulose membranes using the semi-dry turbo transfer system (Biorad). One hour incubation in 5% NFDM in tris-buffered saline .1% tween20 (TBST) was used for blocking, followed by overnight incubation at 4°C with primary antibody diluted in 5% NFDM TBST. anti-TMEFF2 (Sigma HPA015587) 1:1000, anti-PSA (abcam 76113) 1:1000, anti-FKBP5 (abcam 2901) 1:1000, anti-AR (Cell Signaling D6F11) 1:1000, anti-Caspase 3 (Cell Signaling Technology 9662S) 1:1000, anti-H2AX (Cell Signaling Technology 2595S) 1:1000, anti-phospho-H2AX Ser139 (Cell Signaling Technology 9718S) 1:1000, anti-Calnexin (abcam 22595) 1:4000. Goat anti-rabbit HRP conjugated secondary antibody (ThermoFisher 31460) or Goat anti-mouse HRP conjugated secondary antibody (Santa Cruz sc-2005) were diluted in 5% NFDM at a concentration of 80 ng/ml. Clarity Western ECL (Biorad) or SuperSignal West Femto (Thermo) were used for chemiluminescent detection.

### Quantitative RT PCR

RNA samples were reverse transcribed using iScript reverse transcription supermix (500ng/rxn) (Biorad). RT qPCR reactions were carried out in 96 well plates (25 ng cDNA/well and 200 nM per primer/well) using Sso Advanced Universal SYBR Green supermix (Biorad) and the Biorad CFX96 touch RTPCR detection system. Each reaction (sample/primer set combination) was run in duplicate to ensure accurate loading. Relative gene expression was calculated via ΔΔCT method (Livak & Schmittgen, 2001). 4 housekeeping genes (RPL8, RPL38, PSMA1, PPP2CA) were used for normalization per run, with the geometric mean of CT values being used for normalization of gene expression (Vandesompele et al., 2002). The following primers were used:

TMEFF2: For (AGTGCAACAATGACTATGTGCC), Rev (GATCCTGATCCTGCATCTGTG)
KLK3: For (CTTACCACCTGCACCCGGAG), Rev (TGCAGCACCAATCCACGTCA)
KLK2: For (AGAGGAGTTCTTGCGCCCC), Rev (CCCAGCACACAACATGAACTCT)
TMPRSS2: For (CCTCTGACTTTCAACGACCTAGTG), Rev (TCTTCCCTTTCTCCTCGGTGG)
NKX3-1: For (CAGAGACCGAGCCAGAAAGG), Rev (ACTCGATCACCTGAGTGTGGG)
AR: For (CCAGGGACCATGTTTTGCC), Rev (CGAAGACGACAAGATGGACAA)
EHMT2: For (GGAGCCACCGAGAGAGTTCATG), Rev (ACCAACAGTGACAGTGACAGAGG)
MED1: For (GGAGAATCCTGTGAGCTGTCCG), Rev (ATCTTGTTCTAAGGATTGGAGAGCC)
MED21: For (GCTAACCCTACAGAAGAGTATGCC), Rev (CCTCGATAAACAACATCCTCCAGAC)
PIAS1: For (GGGTTTCTCTACTATGTCCACTTGG), Rev (TATGGAGCCTTCTTATCACAGACAG)
RANBP9: For (GGGTGCACTACAAAGGTCATGG), Rev (ACCTGGTAGTCTATTCATGTTCACAC)
RANBP10: For (GTAACCAGGAGACCAGCGACAG), Rev (GGACTCATCCGTCTGCAGGTC)
SMARCD1: For (TGCTACTCTAGACAACAAGATCCATG), Rev (GGTTACCCACCACATCAGTCATTG)
TAF1: For (CAGAAAAGCAGGTAACACAGGAAGG), Rev (TCACTTCCACTTTCACTCAGCTGG)
USP12: For (CAAACAGGAAGCACACAAACGGATG), Rev (CAACAAGGTCGTACATTCTGTCTGG)
ZMIZ1: For (CCAGACGCTGATGTGGAGGTC), Rev (GGCTTGTGGGAGGTCTTGTTGTC)
PSMA1: For (CTGCCTGTGTCTCGTCTTGTATC), Rev (GGCCCATATCATCATAACCAGCA)
RPL8: For (CACCGTTATCTCCCACAACCCT), Rev (AGCCACCACACCAACCACAG)
RPL38: For (ACTTCCTGCTCACAGCCCGA), Rev (TCAGTTCCTTCACTGCCAAACCG)
PPP2CA: For (TTGATCGCCTACAAGAAAGTTCCC), Rev (CATGGCACCAGTTATATCCCTCC)

Statistics: Two-tailed T tests were used to calculate significant differential gene expression. N=3 or 4. P<.05 was considered to be statistically significant.

### RNA sequencing

Deep paired-end RNA sequencing analysis targeted to the TMEFF2 locus for de novo isoform detection was carried out by Oklahoma Medical Research Foundation’s Sequencing Facility. One sample of LNCaP RNA was used for analysis, and over 317 million reads were obtained after decontamination. The genomic region of focus was the TMEFF2 locus −/+ 1Mbp upstream and downstream (chr2: 190949046 – 193194933). StringTie (v.1.2.3) was used for transcript reconstruction.

For differential gene expression analyses, two RNA-seq experiments were conducted:

RNA-seq #1: LNCaP cells expressing shScramble, shTMEFF2-3, shTMEFF2-4, shTMEFF2-9 or shL3. RNA was extracted 55 hours after transductions.
RNA-seq #2: LNCaP cells expressing shScramble or shTMEFF2-3. Cells were grown in RPMI containing 10% charcoal-stripped FBS (hormone depleted media) for 24 hours beginning at 24 hours after transductions. Cells were then grown in RPMI containing 10% charcoal-stripped FBS with the addition of 10 nM DHT or .0001% EtOH for 24 hours prior to RNA extraction.

In both cases RNA samples were prepared as described in the RNA extractions and lysate preparations section of the materials and methods. Three repeats were analyzed for each RNA sequencing analysis. RNA sequencing and initial statistical analyses were carried out by Novogene. Briefly, RNA integrity was analyzed by Agilent 2100 to ensure sample quality. mRNAs were isolated using polyT magnetic beads, which was followed by fragmentation. Two cDNA libraries were synthesized using random hexamer primers; one with dTTP to dUTP substitutions in the second strand to allow for strand specificity, and one library without substitution. cDNA fragments from both libraries were ligated to NEBNext Adaptor and purified using AMPure XP beads after PCR amplification. 20 million reads were conducted for each sample using an Illumina Next-Generation sequencer. For quality control purposes, error rate distributions and G/C content were analyzed in reads, and low quality reads containing adapter sequences, >10% unknown nucleotides or low Q-score values were eliminated by FastQC (Novogene). STAR was used for mapping clean reads to the human transcriptome and genome, and differential gene expression was determined using the DESeq2 R package. P-values were adjusted using the Benjamini and Hochberg method. Transcripts with average log2 fold change >.5 (n=3) and adjusted p-value < ±.05 were considered significantly differentially expressed.

### RNA seq enrichment analyses

For RNA-seq #1, GSEA software version 4.0.1 was used for gene set enrichment analyses (GSEAs) (Mootha et al., 2003; Subramanian et al., 2005). The following Molecular Signature Database (MSigDB) collections were used for gene sets: Hallmark (50 gene sets), Curated (5,529 gene sets), Regulatory target (3,735 gene sets), GO (10,192 gene sets), Oncogenic (189 gene sets) (www.gsea-msigdb.org). For each comparison, genes were rank ordered according to fold change, from the most upregulated to most downregulated, relative to shScramble expressing cells, and duplicate genes were eliminated. One thousand gene permutations were used during enrichment analyses, and FDR q-values less than .25 were considered significant and less than .05 were considered highly significant.

For RNA-seq #2, significantly differentially expressed genes (log2 fold change > .5, adjusted p value < .05) were determined in response to DHT for shScramble and shTMEFF2-3 expressing cells (shScramble +DHT vs shScramble –DHT; shTMEFF2-3 +DHT vs shTMEFF2-3 –DHT), and in shTMEFF2-3 relative to shScramble expressing cells in the presence and absence of DHT (shTMEFF2-3 +DHT vs shScramble +DHT; shTMEFF2-3 –DHT vs shScramble –DHT). GSEAs were conducted as described for RNA-seq #1.

Essential gene list was obtained from Fei et al 2017 (Fei et al., 2017). All of the top 999 essential genes identified in LNCaP cells were used for the essential gene list. In addition, all 274 AR corregulators from DePriest et al (DePriest et al., 2016) were used for AR coregulatory gene list.

P-values for enrichment analyses were calculated by hypergeometric distribution:

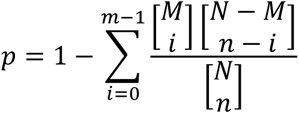

where p = p-value, N = number of total genes, M = number of genes in pathway/gene list, n = number of differentially expressed genes, and i = number of overlapped genes of M and n.

### cWords analyses

The cWords webserver (servers.binf.ku.dk/cwords/ (Rasmussen et al., 2013)) was used to identify 3’ UTR and CDS gene sequences of 6, 7 and 8 nucleotides in length associated with gene downregulation in RNA-seq analyses. Genes were rank ordered from the most downregulated to upregulated for each comparison, and ensembl release 99 3’ UTR and CDS gene sequences were used.

### shRNA seed match analyses

Essential and AR coregulatory gene lists were stratified by downregulation (yes or no), and by the presence of single, two or three or more 6 and 7 nucleotide 3’ UTR sequences found to be most enriched in downregulated genes according to cWords analyses. Ensembl 99 MANE Select transcript 3’ UTR and/or APPRIS annotated 3’UTR sequences were used. 3’UTR sequences were uploaded into R Studio using the seankross/warppipe R package (https://rdrr.io/github/seankross/warppipe/), and 3’UTR sequence length and sequences with specified 6mer and 7mer sequences were identified. Statistics: Pearson’s Chi-square test of independence was used to identify significant associations between 6mer and 7mer 3’ UTR sequences and gene downregulation. P < .05 was considered statistically significant.

### Mouse Xenografts

Animal studies were approved and conducted in accordance with the Institutional Animal Care and Use Committee (IACUC) of the Oklahoma Health Science Center IACUC (animal protocol #17-053-SSHCILA). Mice were housed in groups of 2 or 3 animals per cage. NOD.Cg-Prkdc^scid^ II2rg^tm1 Wjl^/SzJ (NSG) mice (Jackson Labs, Bar Harbor, ME) were used for this study.1.8×10^6^ 22Rv1 cells stably transduced with doxycycline-inducible shScramble or shTMEFF2-9 shRNAs and mixed with Basement membrane extract (R&D systems, Minneapolis, MN) at a 1:1 ratio, were injected subcutaneously into the flanks of mice pre-fed for two days with chow containing doxycycline (Bio-Serv, Flemington, NJ). Mice were maintained on the doxycycline chow diet and tumor growth was monitored using the Biopticon TumorImager. Mice were sacrificed approximately five weeks after injections, and tumors were excised and dehydrated tumor weights were determined. Differences in shScramble and shTMEFF2-9 tumor weight were analyzed statistically by t-test. N = 7.

## ACKNOWLEDGEMENTS

We are grateful to Dr. S. Hayward (NorthShore Research Institute) for sharing cell lines. The authors acknowledge Cody Bullock’s help with mycoplasma detection. Research reported in this publication was supported in part by the National Cancer Institute Cancer Center Support Grant P30CA225520, COBRE P20GM103639 and the Oklahoma Tobacco Settlement Endowment Trust contract awarded to the University of Oklahoma Stephenson Cancer Center and used the Tissue Pathology and the Molecular Biology and Cytometry Research Shared Resources. The content is solely the responsibility of the authors and does not necessarily represent the official views of the National Institutes of Health or the Oklahoma Tobacco Settlement Endowment Trust.

---

**Figure 1 – figure supplement 1.**
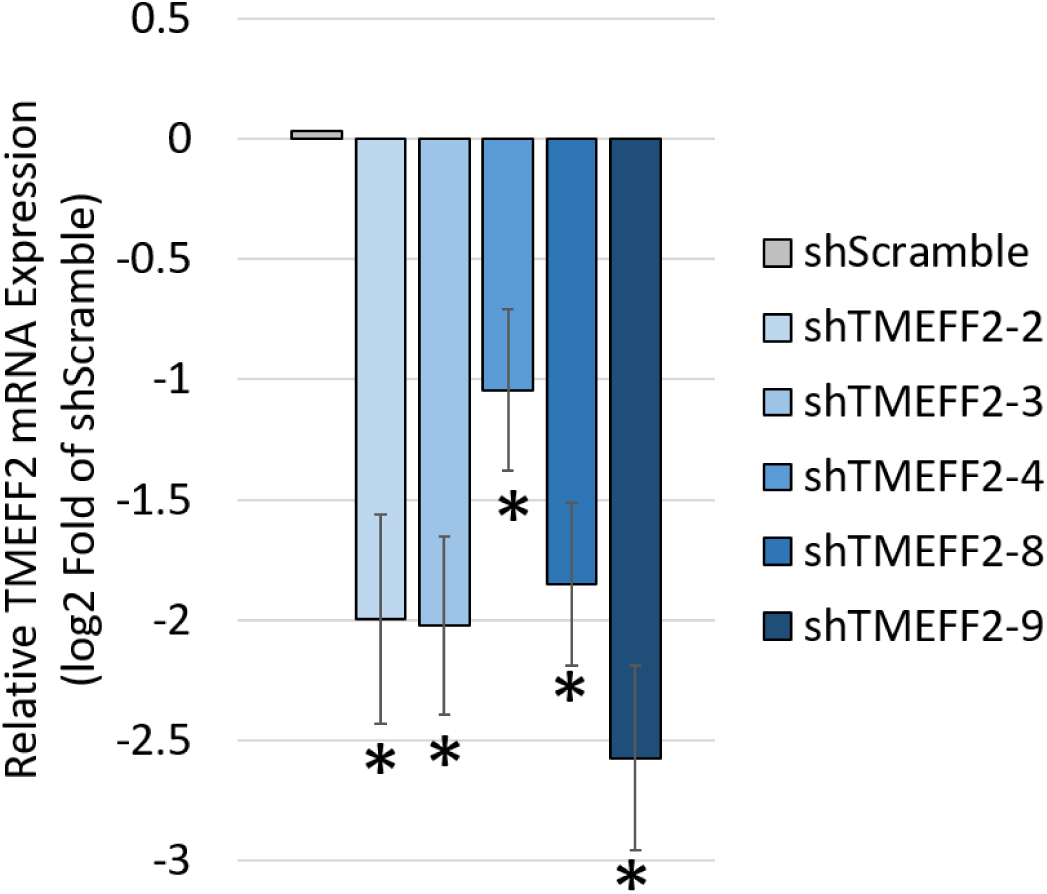
TMEFF2 knockdown in response to TMEFF2-targeted shRNA. TMEFF2 mRNA expression in LNCaP cells expressing shTMEFF2-2, shTMEFF2-3, shTMEFF2-4, shTMEFF2-8 or shTMEFF2-9 relative to cells expressing shScramble control. RNA was extracted 72 hours after transductions, and mRNA expression was determined by RT qPCR. N=3, error bars ±SD, * p<.05 determined by t-test.

**Figure 1 – figure supplement 2.**
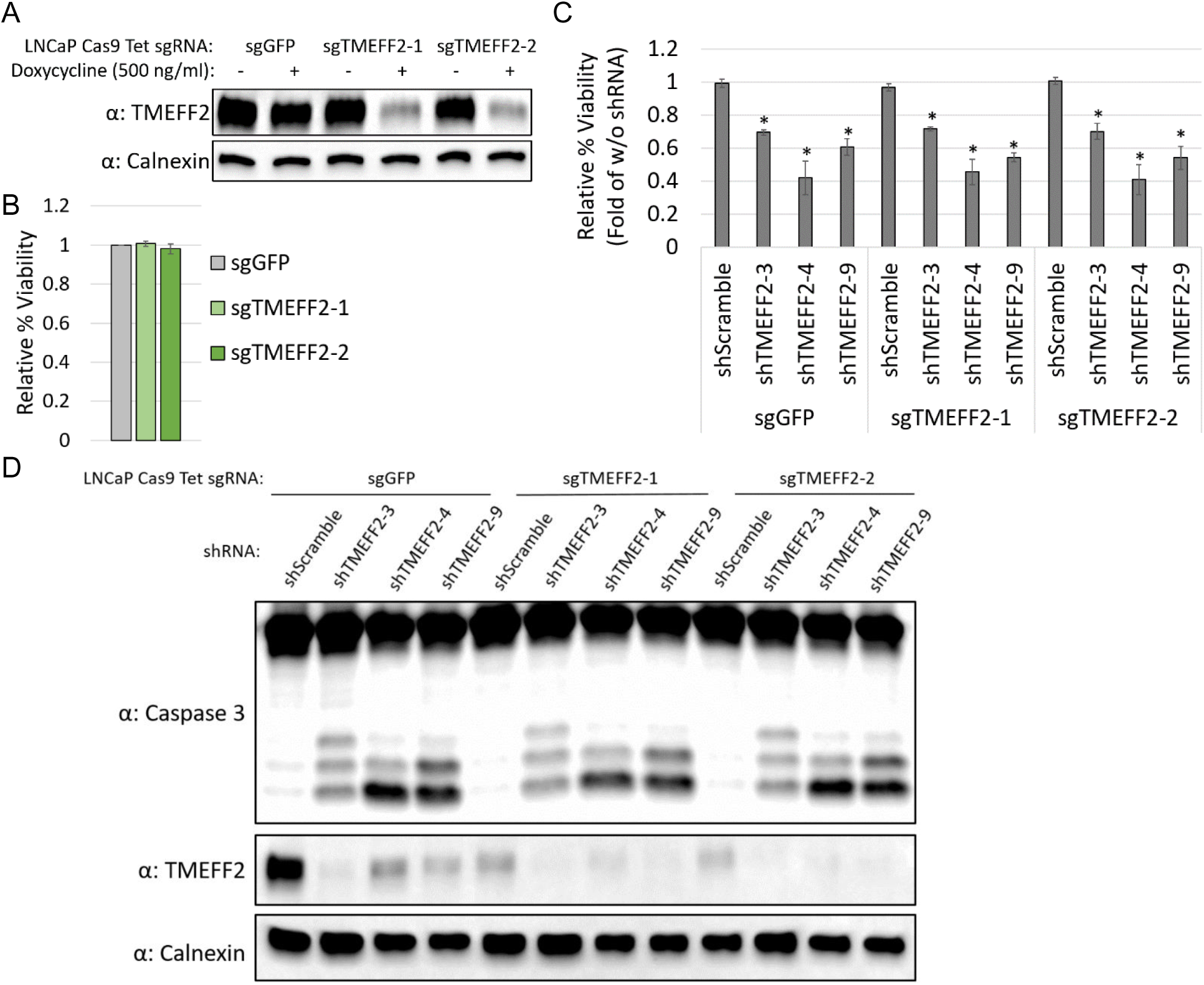
Loss of viability in response to TMEFF2-targeted shRNA is independent of TMEFF2 protein levels. (**A**) Western blot analysis showing doxycycline-induced TMEFF2 knockdown in LNCaP Cas9 cells expressing doxycycline-inducible TMEFF2-targeted sgRNAs (sgTMEFF2-1 and sgTMEFF2-2) after cells were grown for 10 days in the presence and absence of 500 ng/ml doxycycline. Doxycycline-inducible GFP targeted sgRNA (sgGFP) served as a negative control. Calnexin was used as a loading control. (**B**) Percent viability of LNCaP Cas9 sgTMEFF2-1 and sgTMEFF2-2 cells lines relative to LNCaP Cas9 sgGFP cell line. Viability determined by trypan blue after cells were grown in 500 ng/ml doxycycline for 10 days to induce sgRNA expression. N=3, error bars ±SD. (**C**) Relative percent viability and (**D**) western blot analysis showing TMEFF2 expression and caspase 3 cleavage in lysates from LNCaP Cas9 sgGFP, sgTMEFF2-1 and sgTMEFF2-2 cell lines grown in 500 ng/ml dox and subsequently transduced with plasmids expressing TMEFF2-targeted shRNAs or shScramble control. Viability was measured by trypan blue 96 hours after shRNA transductions. Viability is presented as percent viability relative to the viability of cells without shRNA expression. N=3, error bars ±SD, * p<.05 determined by t-test. Lysates were also obtained 96 hours after shRNA transductions, and calnexin was used as a loading control.

**Figure 1 – figure supplement 3.**
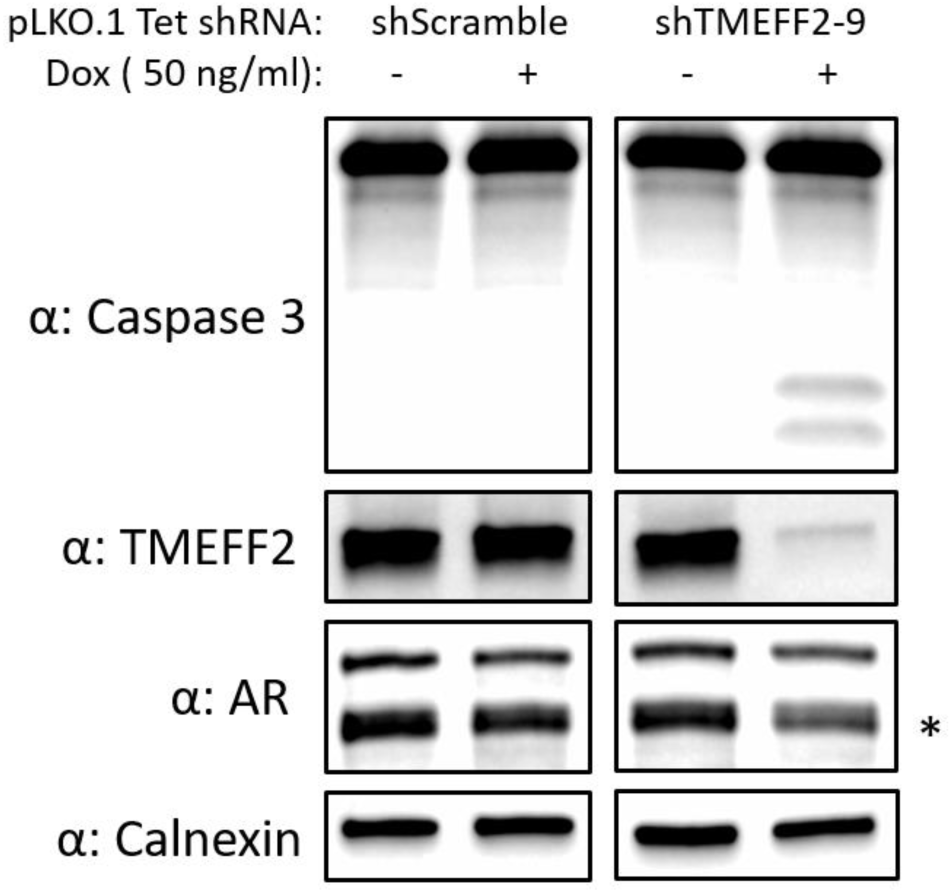
Doxycycline inducible TMEFF2-targeted shRNA expression induces caspase 3 cleavage and reduces AR protein expression in 22Rv1 cells. Western blot analysis of Caspase 3 cleavage, TMEFF2 and AR protein (* AR-V7) expression in lysates from 22Rv1 shScramble and shTMEFF2-9 cell lines grown in the presence and absence of 50 ng/ml doxycycline for 5 days to induce shRNA expression. Calnexin was used as a loading control.

**Figure 1 – figure supplement 4.**
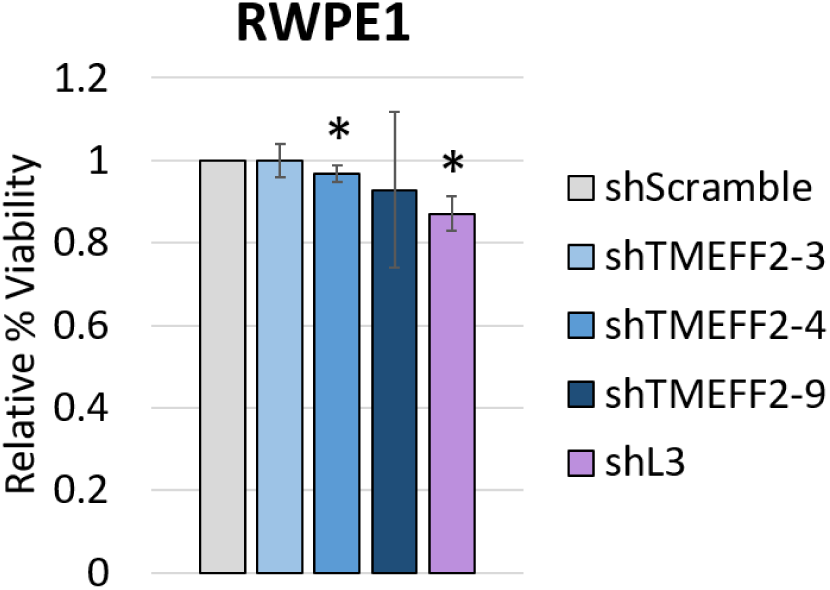
TMEFF2-targeted shRNAs and shL3 have minimal effect on normal prostate cell viability. Percent viability of RWPE1 cells expressing shTMEFF2-3, shTMEFF2-4, shTMEFF2-9 or shL3 relative to the viability of cells expressing shScramble control shRNA. Viability was measured by trypan blue 120 hours after transductions. N=3, error bars ±SD, * p<.05 determined by t-test.

**Figure 2 – figure supplement 1.**
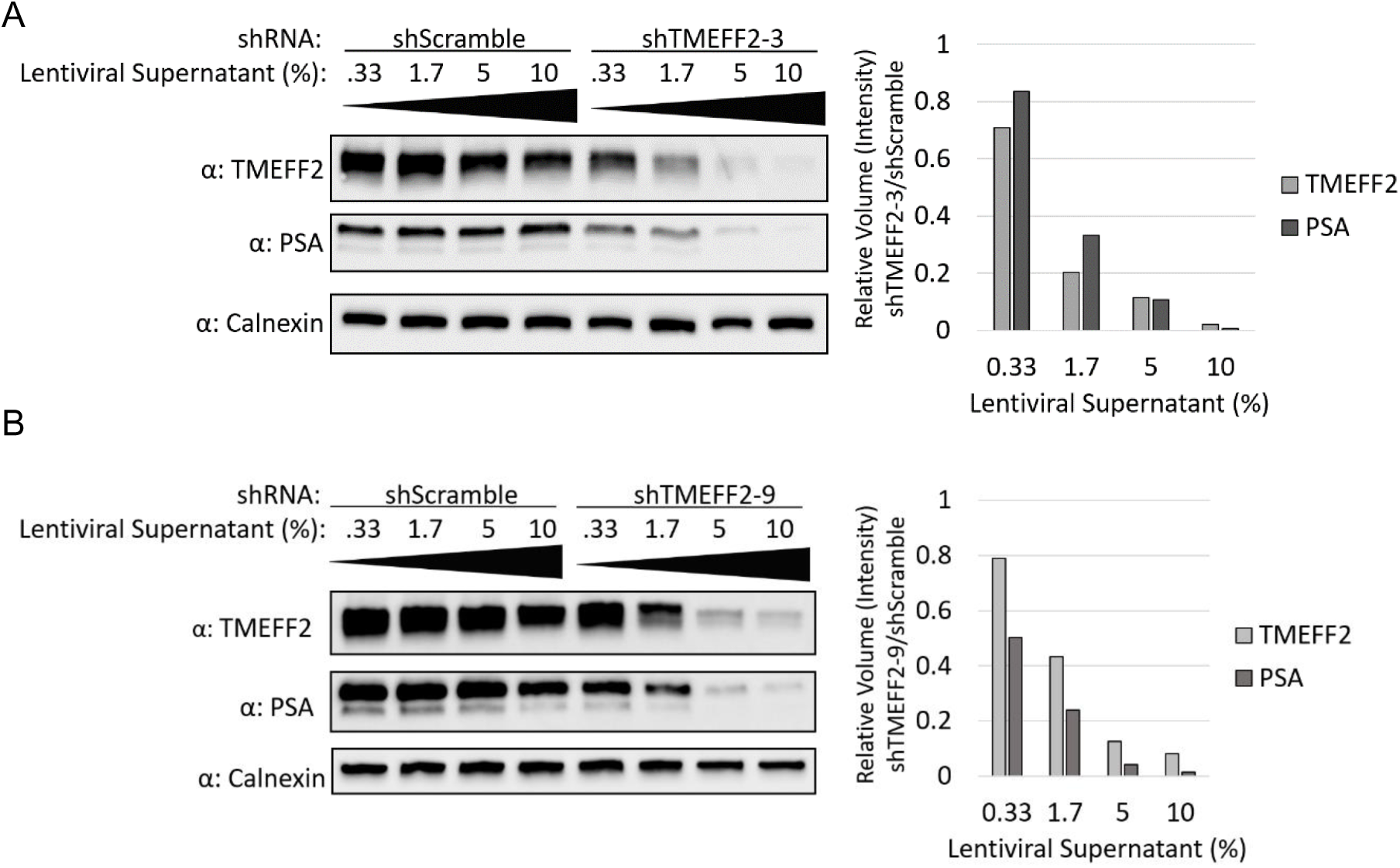
Low TMEFF2-targeted shRNA expression reduces PSA protein levels. Western blot analysis showing PSA and TMEFF2 protein levels in response to increasing dose of lentiviral particles containing plasmids expressing shTMEFF2-3 (**A**) or shTMEFF2-9 (**B**) and shScramble shRNA in LNCaP cells, four days after transduction. Dose is presented as percentage of lentiviral supernatant in transduction media. See right panel for graphical representation of Calnexin normalized PSA and TMEFF2 levels in response to shTMEFF2-3 expression relative to shScramble control at each dose. Band intensity was quantified using Biorad Image Lab.

**Figure 2 – figure supplement 2.**
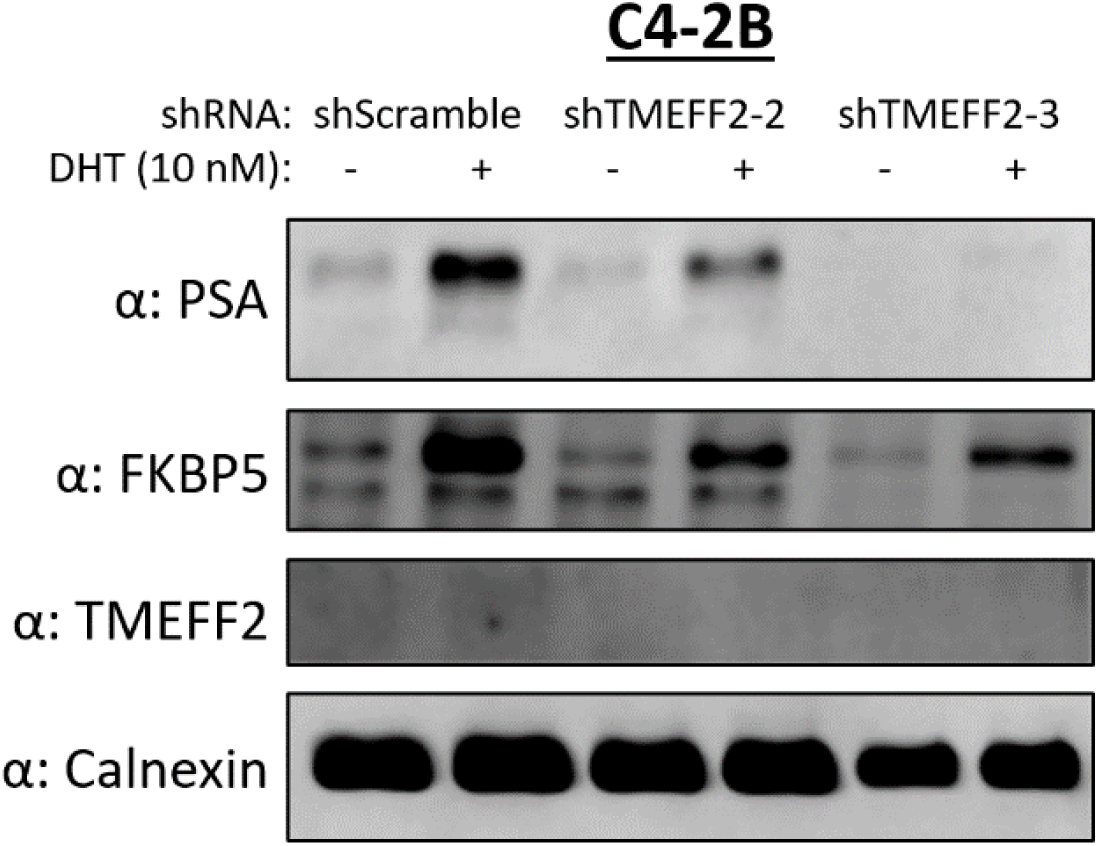
TMEFF2-targeted shRNAs reduce androgen responsive protein levels. TMEFF2 and androgen responsive protein expression in lysates from C4-2B cells expressing shTMEFF2-2, shTMEFF2-3 or shScramble control shRNAs and grown in the presence and absence of 10 nM DHT. Calnexin was used as a loading control for all western blots.

**Figure 2 – figure supplement 3.**
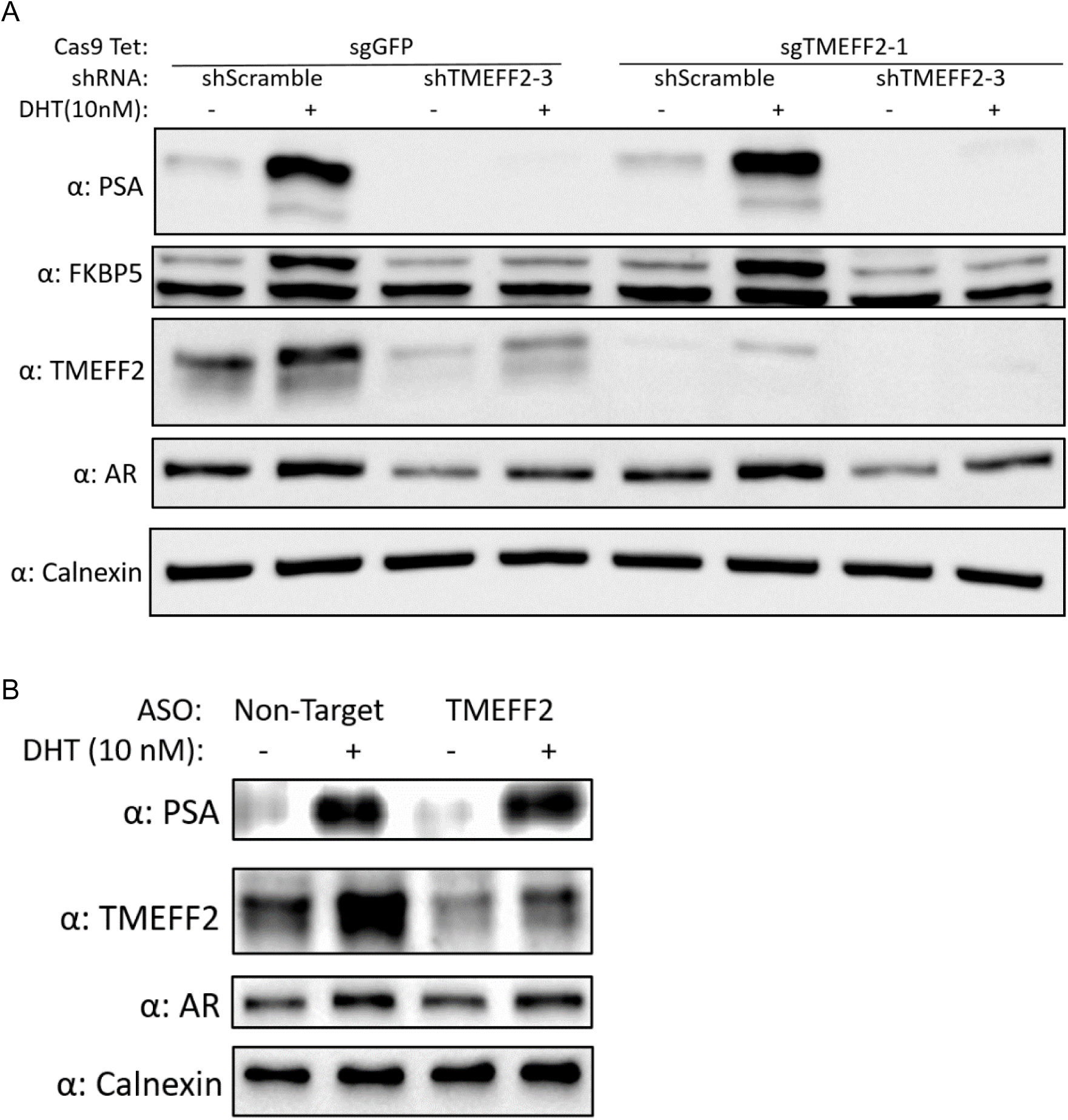
Inhibition of androgen signaling in response to TMEFF2-targeted shRNA is independent of TMEFF2 protein levels. Western blot analysis showing TMEFF2, AR, PSA and FKBP5 protein levels. Lysates were from LNCaP Cas9 sgGFP and sgTMEFF2-1 cell lines, grown for 2 weeks in 500 ng/ml doxycycline to induce TMEFF2 knockdown, followed by 2 weeks in the absence of doxycycline, then transduced with plasmids expressing shScramble or shTMEFF2-3 shRNAs, and grown in the presence or absence of 10 nM DHT. Western blot analysis showing PSA, TMEFF2 and AR protein levels in lysates from LNCaP cells transfected with Non-Target or a pool of TMEFF2 targeting ASO’s, and grown in the presence and absence of DHT. Calnexin was used as a loading control for all western blots.

**Figure 2 – figure supplement 4.**
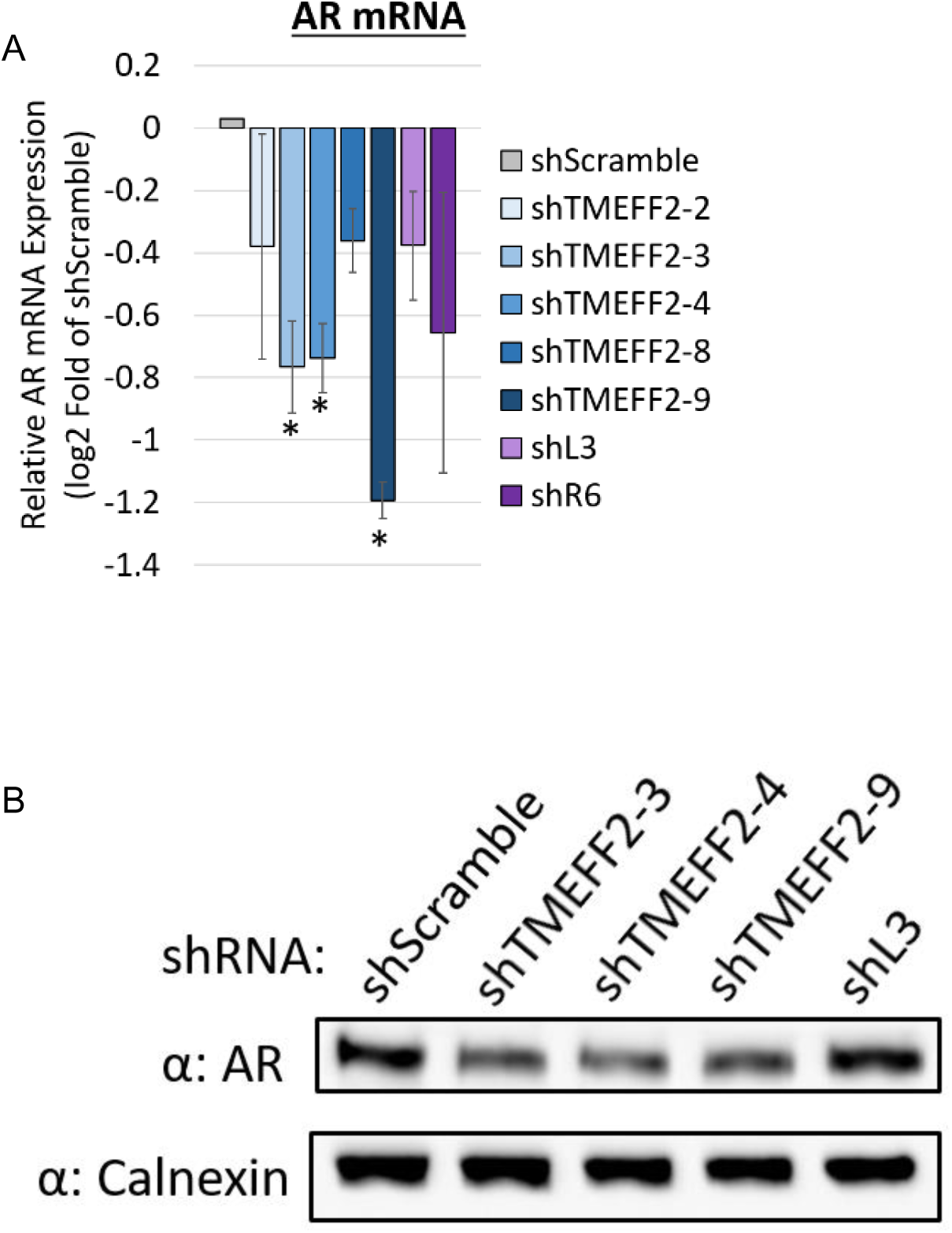
TMEFF2-targeted shRNAs reduce AR protein levels, while shL3 does not. (**A**) AR mRNA expression in LNCaP cells expressing TMEFF2-targeted shRNAs, shL3 or shR6 relative to cells expressing shScramble control. RNA was extracted 72 hours after transductions, and mRNA expression was determined by RT qPCR. RPL8, RPL38, PSMA1 and PPP2CA were the housekeeping genes used for normalization. N=3, error bars ±SD, * p<.05 determined by t-test. (**B**) Western blot analysis showing AR protein expression in lysates obtained from LNCaP cells expressing shTMEFF2-3, shTMEFF2-4, shTMEFF2-9, shL3 or shScramble control shRNA. Lysates were obtained 96 hours after transduction. Calnexin was used as a loading control.

**Figure 2 – table supplement 1.** Transcripts identified by deep RNA-seq of TMEFF2 locus and the number of shRNAs targeting each transcript. Table shows four TMEFF2 isoforms and ten lncRNAs detected as being expressed from the TMEFF2 locus in LNCaP cells using deep RNA-seq analysis (317 million good reads). A list of detected transcripts is provided along with information regarding location, transcript/gene ID/symbol, length, number of exons, relative transcript abundance (coverage/ Fragments Per Kilobase of transcript per Million mapped reads (FPKM)), and the number of shRNA target sequences out of nine total TMEFF2-targeted shRNAs.

| Chr | strand | Hg38 Start | End | Transcript ID | # exons | Length | Gene ID | Gene Symbol | Coverage | FPKM | # shRNA Target Sequences |
| --- | --- | --- | --- | --- | --- | --- | --- | --- | --- | --- | --- |
| 2 | - | 191950017 | 192195709 | ENST00000272771.9 | 10 | 2604 | ENSG00000144339.11 | TMEFF2 | 605.5575 | 63319.688 | 9 |
| 2 | - | 191949043 | 192194916 | ENST00000392314.5 | 10 | 2842 | ENSG00000144339.11 | TMEFF2 | 263.03003 | 27503.549 | 8 |
| 2 | - | 192176743 | 192194524 | ENST00000409056.3 | 4 | 3364 | ENSG00000144339.11 | TMEFF2 | 65.105675 | 6807.7285 | 2 |
| 2 | + | 192032548 | 192044525 | lnc-OBFC2A-5:20 | 3 | 1152 | lnc-OBFC2A-5 | . | 23.760929 | 2484.5447 | 0 |
| 2 | + | 192035256 | 192037103 | ENST00000625221.1 | 3 | 805 | ENSG00000233766.7 | AC098617.1 | 21.792868 | 2278.7559 | 0 |
| 2 | + | 192021912 | 192029036 | lnc-OBFC2A-5:4 | 3 | 277 | lnc-OBFC2A-5 | . | 21.315186 | 2228.8071 | 0 |
| 2 | + | 192032368 | 192035380 | ENST00000598327.3 | 3 | 638 | ENSG00000233766.7 | AC098617.1 | 18.98439 | 1985.0894 | 0 |
| 2 | - | 191992664 | 191999205 | ENST00000487771.1 | 3 | 278 | ENSG00000144339.11 | TMEFF2 | 12.179338 | 1273.5238 | 1 |
| 2 | + | 192036170 | 192038059 | lnc-OBFC2A-5:23 | 4 | 624 | lnc-OBFC2A-5 | . | 11.182352 | 1169.2748 | 0 |
| 2 | + | 191922546 | 192036829 | ENST00000428980.6 | 5 | 1207 | ENSG00000233766.7 | AC098617.1 | 10.120426 | 1058.2352 | 0 |
| 2 | + | 192035210 | 192037121 | ENST00000602099.3 | 4 | 708 | ENSG00000233766.7 | AC098617.1 | 0.411862 | 43.066055 | 0 |
| 2 | + | 192035210 | 192037121 | lnc-OBFC2A-5:21 | 4 | 708 | lnc-OBFC2A-5 | . | 0.346128 | 36.192627 | 0 |
| 2 | + | 192035256 | 192037103 | lnc-OBFC2A-5:22 | 3 | 805 | lnc-OBFC2A-5 | . | 0.287261 | 30.037245 | 0 |
| 2 | + | 192030605 | 192036747 | ENST00000599681.5 | 5 | 767 | ENSG00000233766.7 | AC098617.1 | 0.002465 | 0.257751 | 0 |

**Figure 3 – figure supplement 1.**
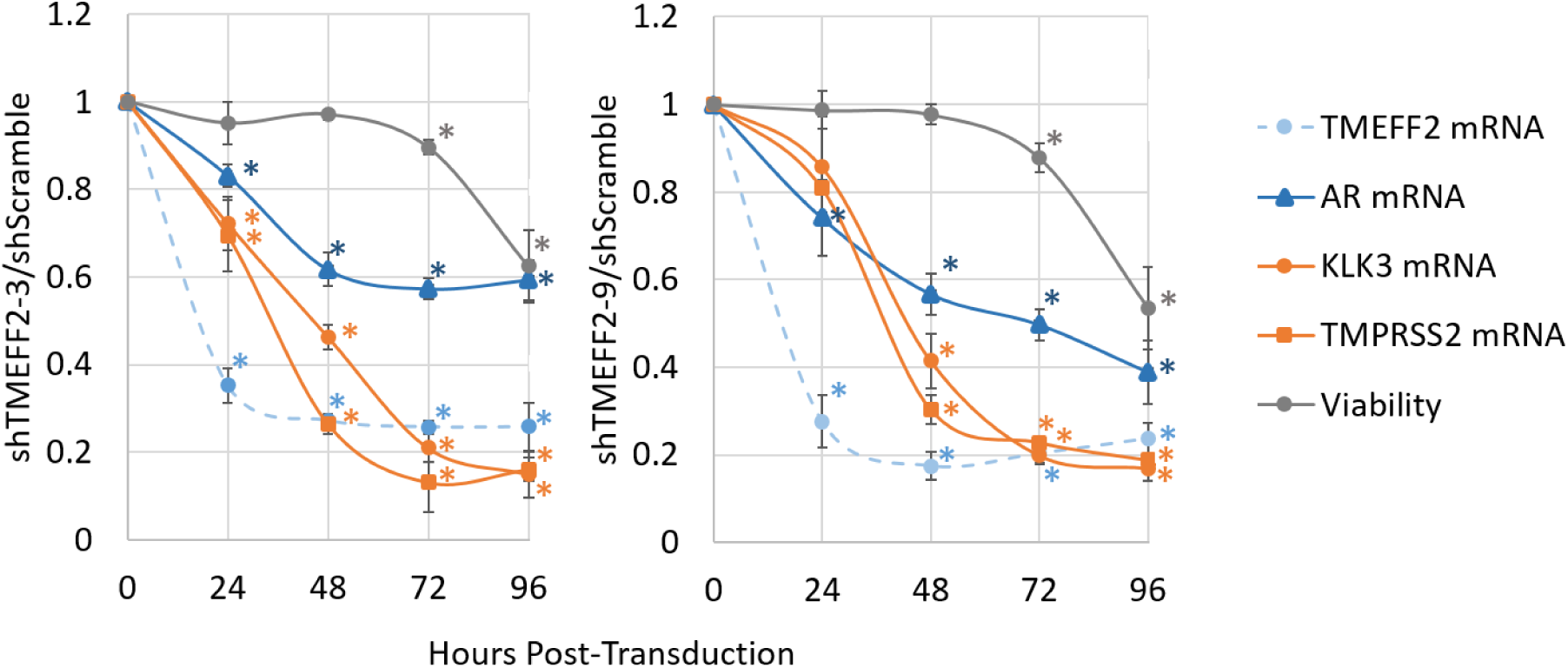
Reductions in AR and androgen responsive genes occur prior to loss of viability in response to TMEFF2-targeted shRNAs. Time course analysis of TMEFF2, KLK3, TMPRSS2 and AR mRNA expression and viability of LNCaP cells expressing shTMEFF2-3 (left panel) or shTMEFF2-9 (right panel) relative to cells expressing shScramble. Viability was determined by trypan blue, and mRNA expression was determined by RT qPCR. RPL8, RPL38, PSMA1 and PPP2CA were the housekeeping genes used for normalization. N=3, error bars ±SD, * p<.05 determined by t-test (for H_0_: viability or expression = 1).

**Figure 3 – figure supplement 2.**
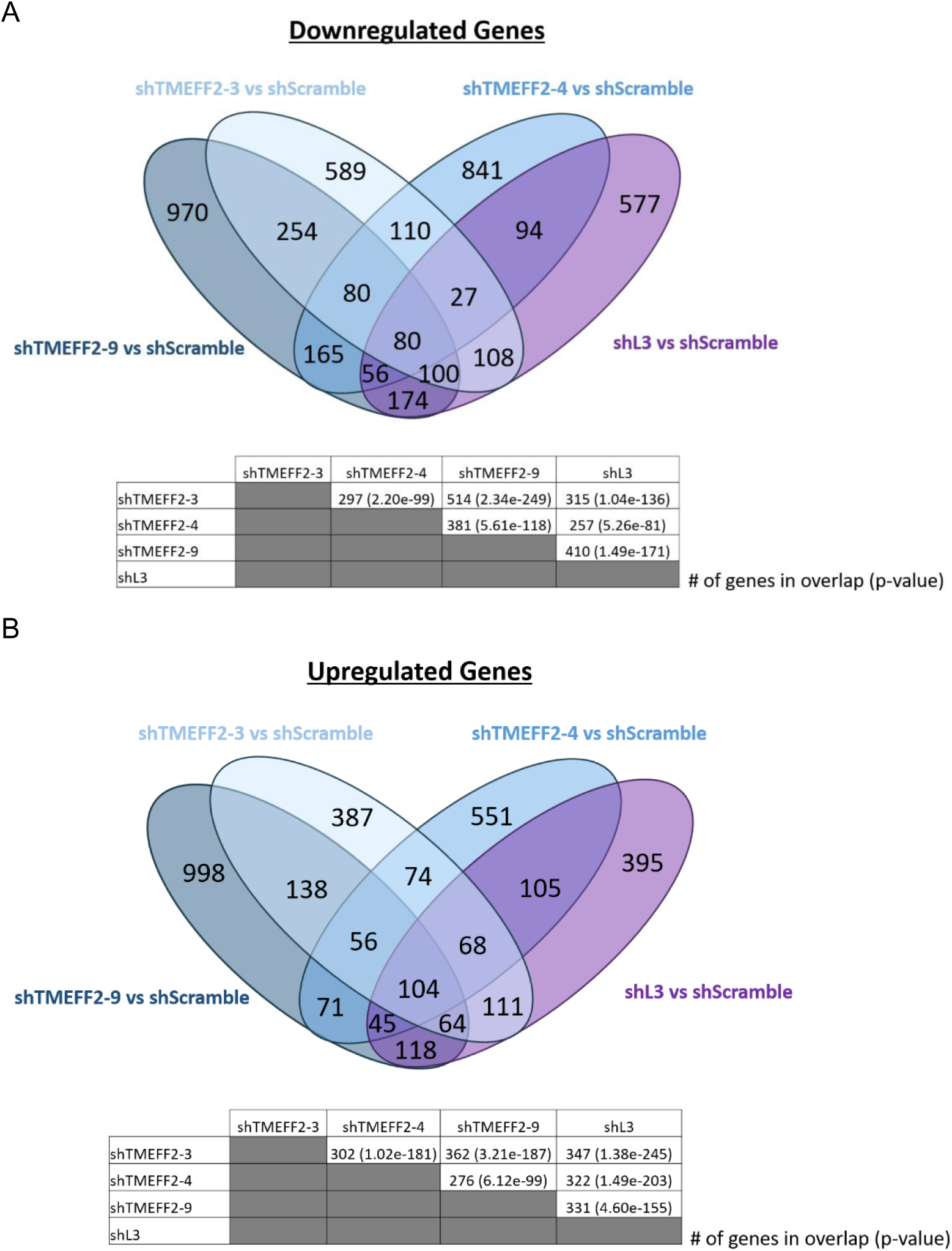
shTMEFF2-3, shTMEFF2-4, shTMEFF2-9 and shL3 expressing LNCaP cells exhibit overlaps in DEGs. Venn diagrams show the number of genes significantly downregulated (**A**) and upregulated (**B**) for each comparison from RNA-seq analysis of LNCaP cells expressing designated shRNAs. Tables show the number of overlapping DEGs and p-values indicating the significance of overlaps for pairwise comparisons of each shRNA based on a hypergeometric distribution.

**Figure 3 – figure supplement 3.**
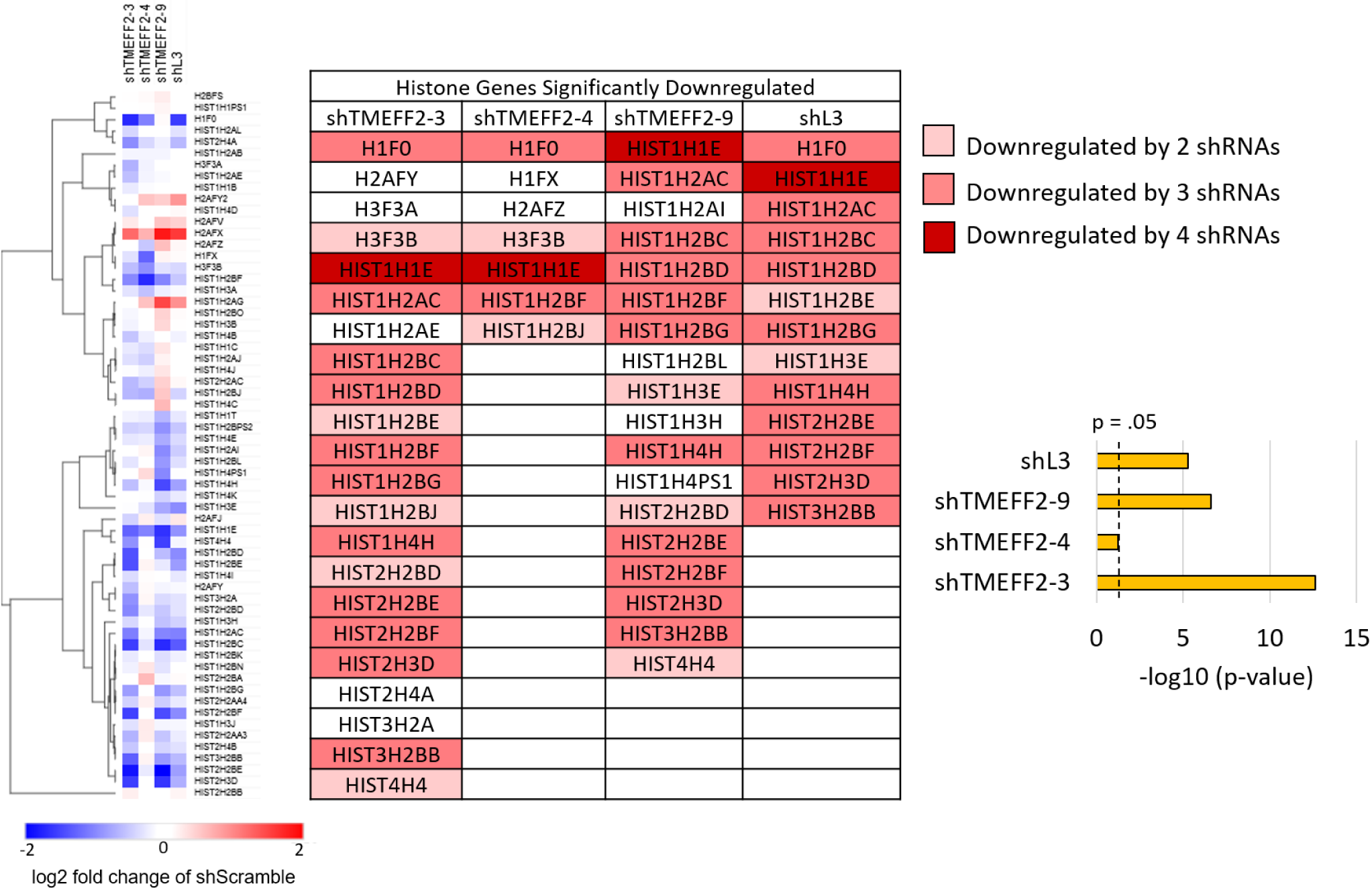
shTMEFF2-3, shTMEFF2-4, shTMEFF2-9 and shL3 downregulate histone mRNAs. Heatmap (Left panel) shows the expression of histone genes in LNCaP cells expressing designated shRNAs relative to LNCaP cells expressing shScramble shRNA according to RNA-seq analysis. Table shows histone genes significantly downregulated by each shRNA relative to LNCaP cells expressing shScramble shRNA. Commonly downregulated histone genes are highlighted in red. Bar graph shows –log10 p-value for enrichment of histone genes among significantly downregulated genes by each shRNA relative to shScramble expressing LNCaP cells. –log10 p-values are based on hypergeometric distribution.

**Figure 3 –figure supplement 4.**
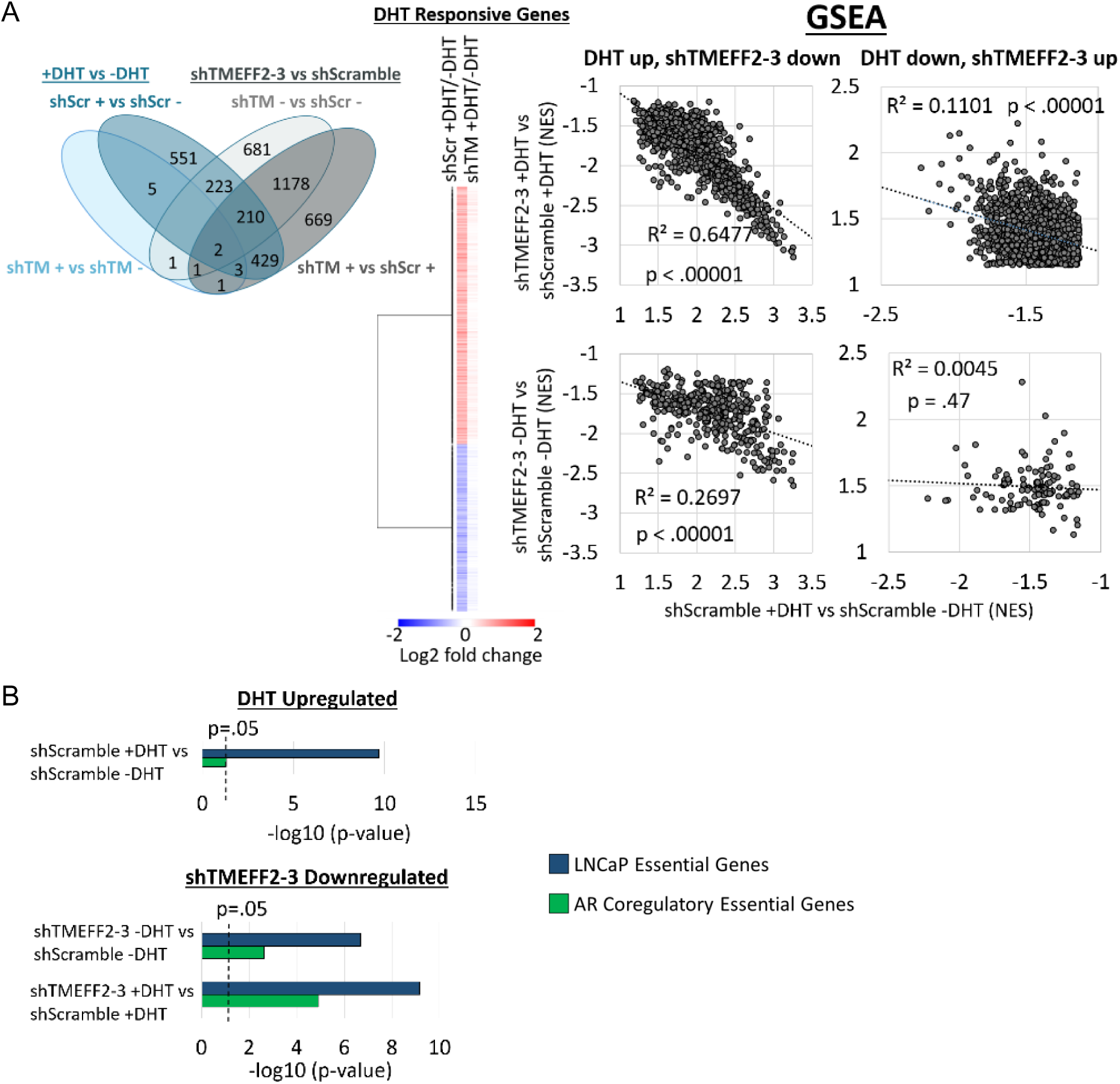
shTMEFF2-3 inhibits global androgen-induced transcriptional response. (**A**) Venn diagram (left panel) shows the number of significantly differentially expressed genes (DEGs) between each comparison (shTM: shTMEFF2-3; Scr: shScramble; −: − DHT; +: + DHT) from RNA-seq analysis of LNCaP cells expressing designated shRNAs and grown in the presence or absence of DHT. Heatmap (middle panel) shows fold change in gene expression of DEGs in the response to DHT in LNCaP cells expressing shTMEFF2-3 or shScramble shRNA. Correlations (right panel) of NES values from GSEA of gene sets that exhibit opposite enrichment with shTMEFF2-3 expression and DHT treatment in shScramble expressing LNCaP cells. (**B**) –log10 p-value for enrichment of AR coregulatory genes (DePriest et al., 2016) and LNCaP essential genes (Fei et al., 2017) among genes significantly upregulated or downregulated by DHT in LNCaP cells expressing shScramble shRNA (Top), and among genes significantly downregulated by shTMEFF2-3 relative to shScramble expressing LNCaP cells (Bottom). –log10 p-values are based on hypergeometric distribution.

**Figure 3 –figure supplement 5.**
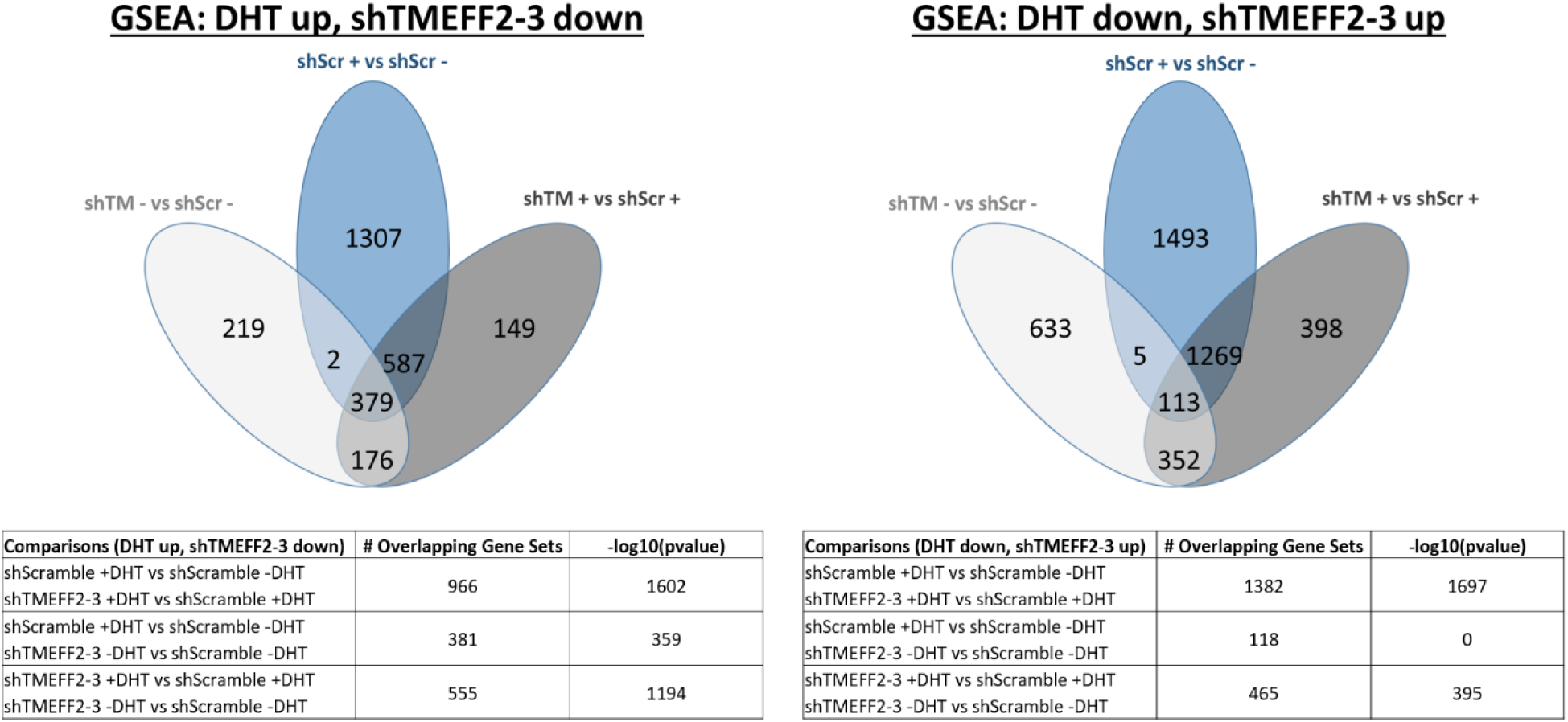
DHT treatment and shTMEFF2-3 oppositely regulate a significant number of gene sets. Venn diagrams show the number of gene sets that are significantly downregulated by shTMEFF2-3 or upregulated by DHT in shScramble expressing LNCaP cells (left panel) and significantly upregulated by shTMEFF2-3 or downregulated by DHT in shScramble expressing LNCaP cells (right panel). The number of overlapping gene sets and the significance of the overlap, –log10(p-values) based on hypergeometric distribution, are shown. (shTM: shTMEFF2-3; shScr: shScramble; −: − DHT; +: + DHT). Enriched gene sets were determined by GSEA (q-value < .25) of RNA-seq data. 19,695 total MsigDB gene sets queried. Top 100 DHT upregulated and downregulated gene sets are listed in Figure 3 – table supplement 2 and 3, respectively.

**Figure 3 – table supplement 1.**
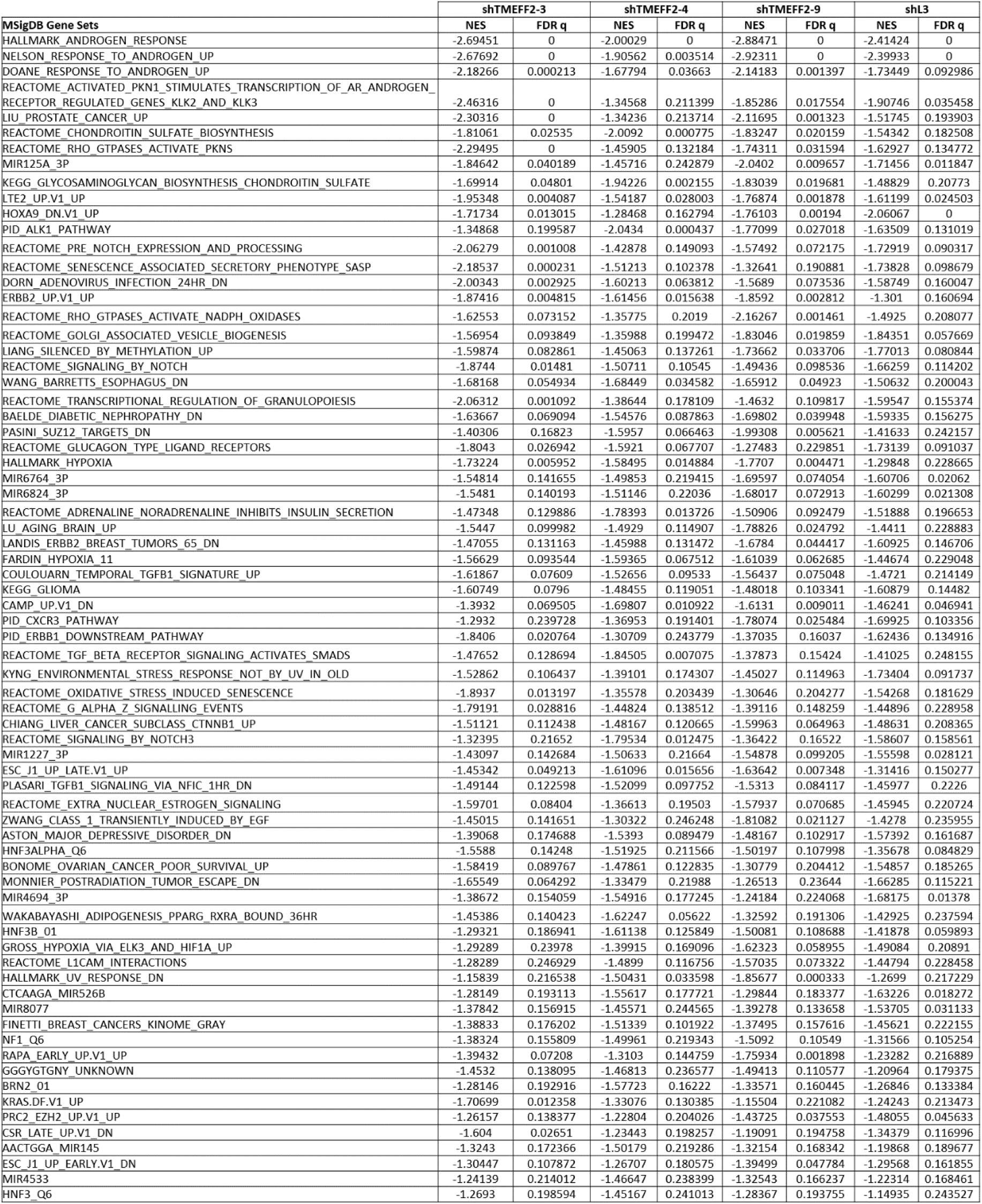
Common enriched gene sets resulting from shTMEFF2-3, shTMEFF2-4, shTMEFF2-9 and shL3 expression in LNCaP cells. Normalized enrichment scores (NES) and q-values for significantly enriched gene sets (GSEA; q-value <.25, for each shRNA) common to LNCaP cells expressing the designated shRNAs compared to cells expressing the shScramble control. q-value=0 : <.001.

**Figure 3 – table supplement 2.**
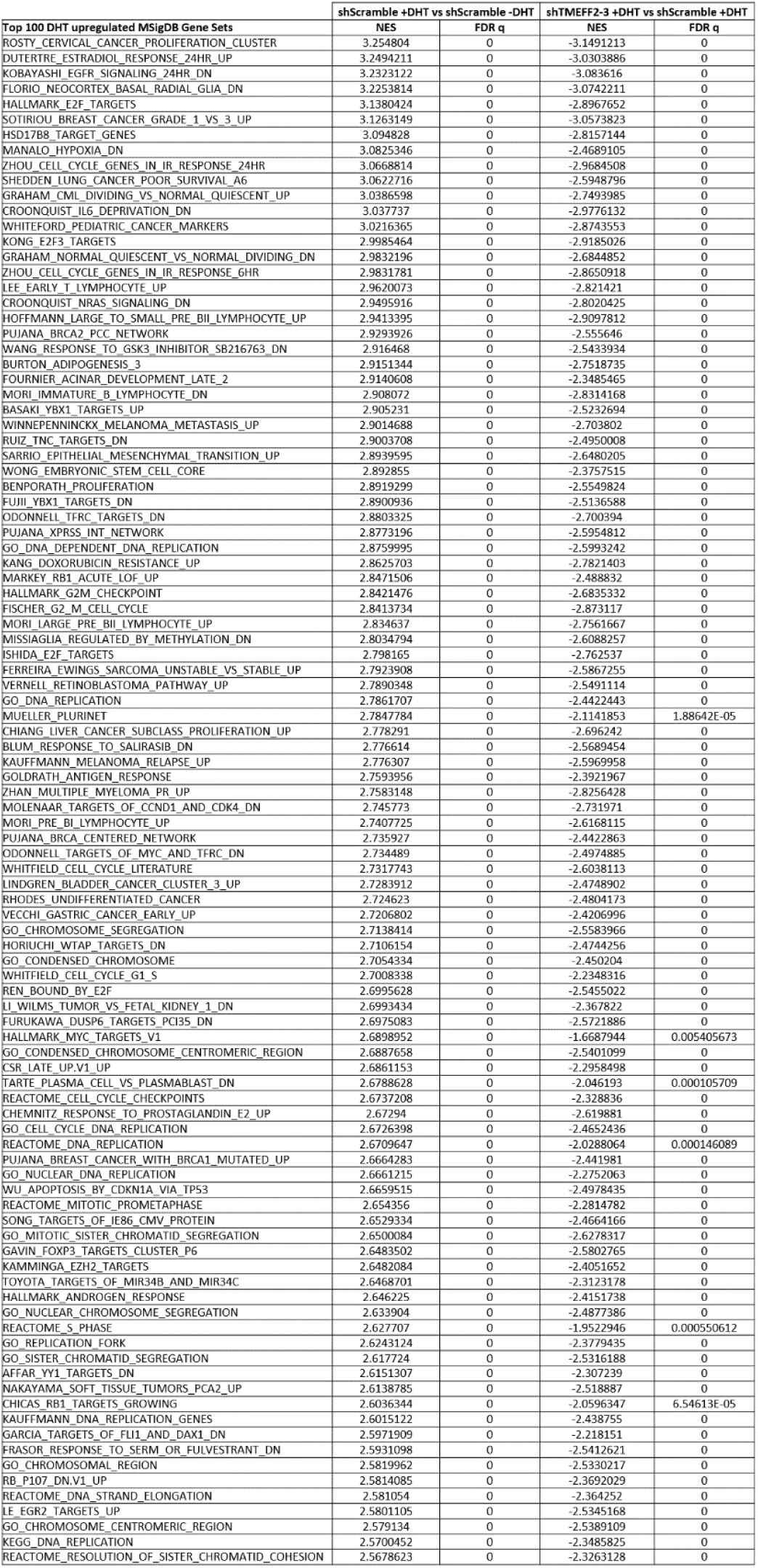
shTMEFF2-3 regulation of the top 100 DHT upregulated gene sets. Table showing the top 100 gene sets (based on GSEA NES value) upregulated by DHT treatment in shScramble LNCaP cells. NES and q-values are shown for gene set regulation by DHT (shScramble +DHT vs shScramble –DHT) and by shTMEFF2-3 in the presence of DHT (shTMEFF2-3 +DHT vs shScramble + DHT). Positive NES values indicate gene set upregulation. Negative NES values indicate gene set downregulation. q-value < .25 indicates significantly enriched gene set. q-value=0 : <.001.

**Figure 3 – table supplement 3.**
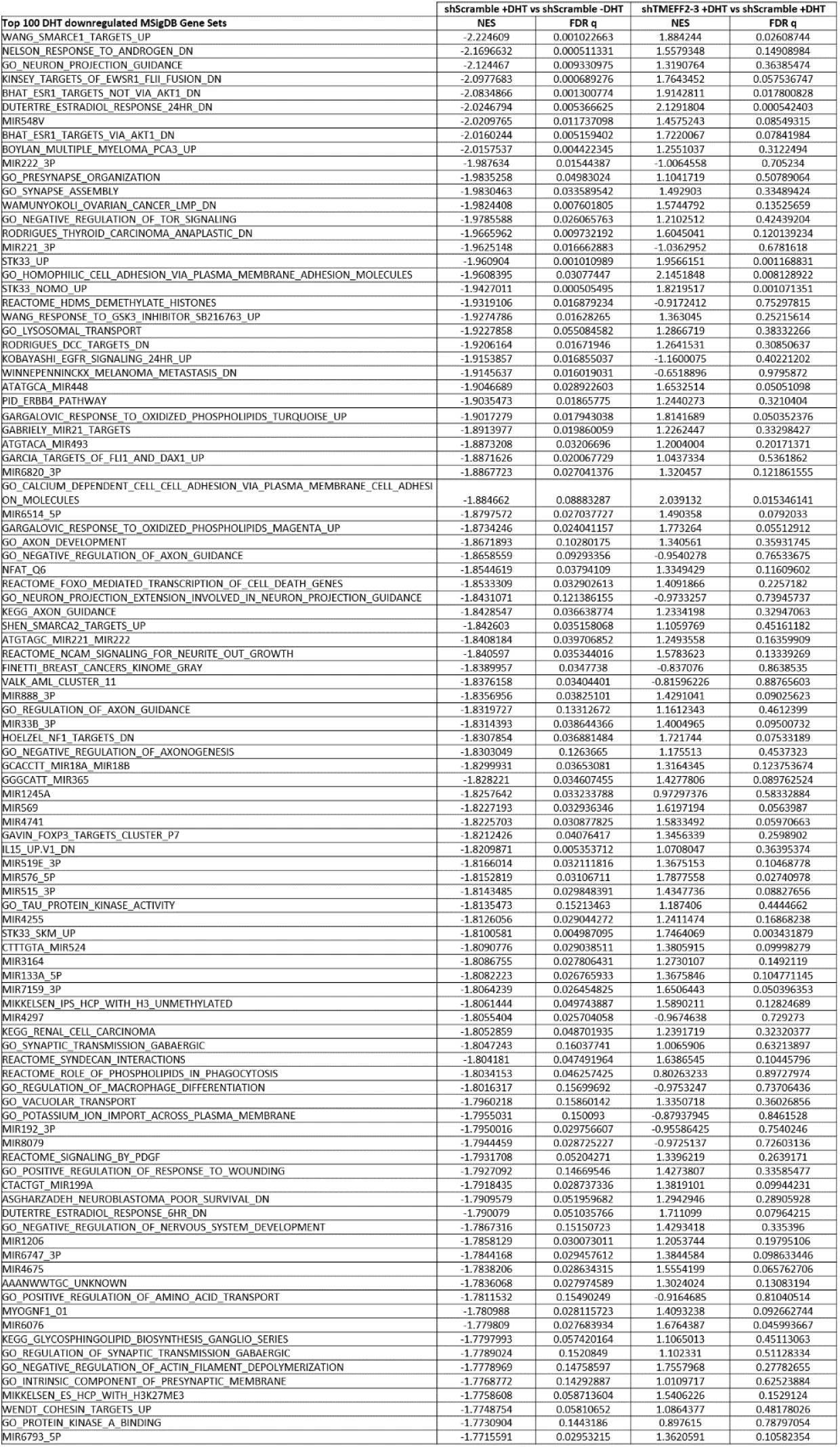
shTMEFF2-3 regulation of the top 100 DHT downregulated gene sets. Table showing the top 100 gene sets (based on GSEA NES value) downregulated by DHT treatment in shScramble LNCaP cells. NES and q-values are shown for gene set regulation by DHT (shScramble +DHT vs shScramble –DHT) and by shTMEFF2-3 in the presence of DHT (shTMEFF2-3 +DHT vs shScramble +DHT). Positive NES values indicate gene set upregulation. Negative NES values indicate gene set downregulation. q-value < .25 indicates significantly enriched gene set.

**Figure 4 – figure supplement 1.**
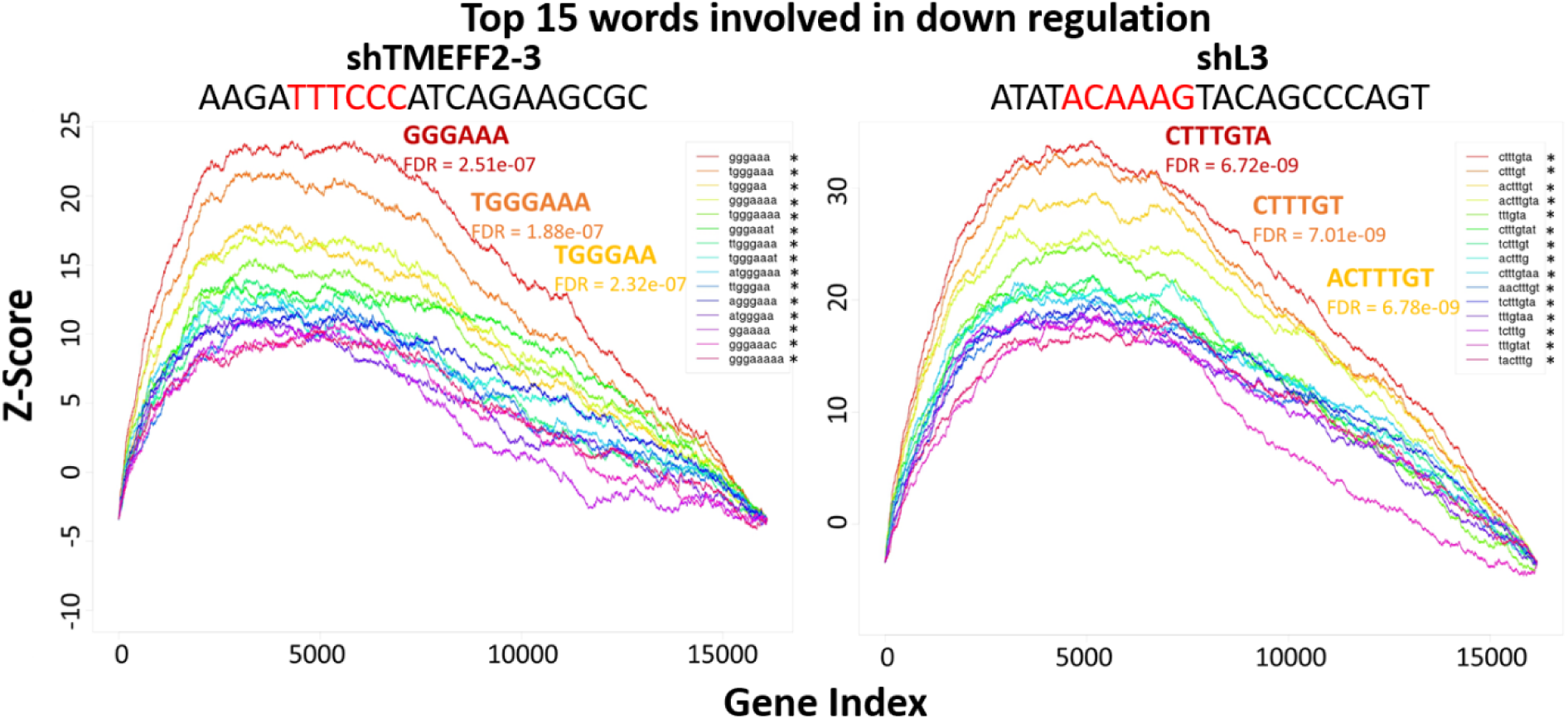
The top 3’ UTR motifs associated with shRNA-mediated gene downregulation are complementary to potential shRNA seed sequences. cWords enrichment plots showing the top 15 enriched 6mer, 7mer and 8mer in the 3’ UTR of genes downregulated by designated shRNAs according to RNA-seq analyses. Y-axis shows Z-score enrichment values. X-axis contains rank ordered genes from the most downregulated to upregulated expression. The top 3 enriched 3’ UTR sequences and FDR values are labeled (red: most enriched, orange: second most enriched, yellow: third most enriched). Unprocessed shRNA guide strand sequences are above each plot, and the potential 6mer seed sequence complementary to the most enriched 6mer 3’ UTR sequence associated with gene downregulation is in red. * indicates enriched sequences that are complementary to potential shRNA seed motifs.

**Figure 4 – figure supplement 2.**
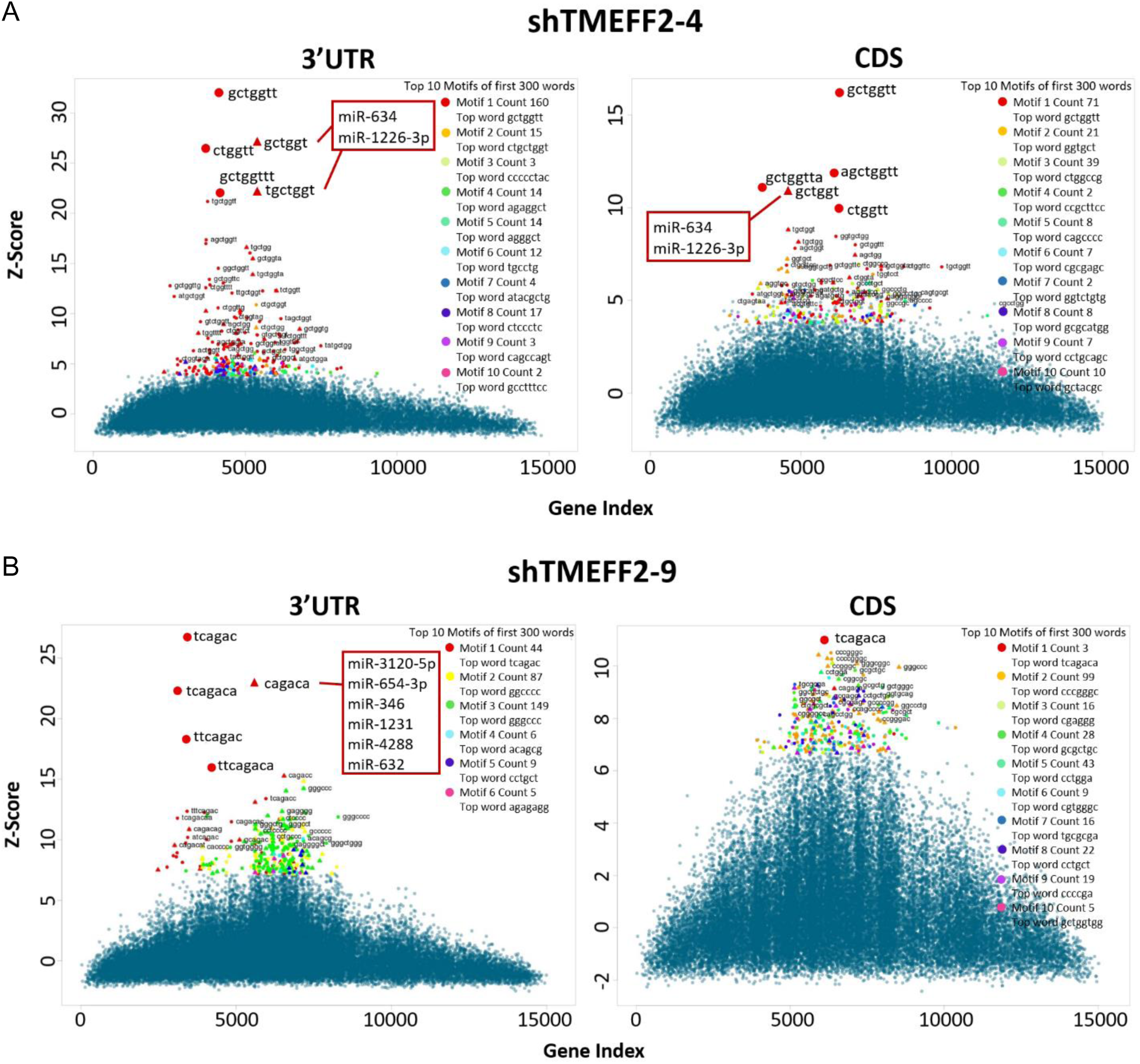
The top 3’ UTR motifs associated with shRNA-mediated gene downregulation are complementary to potential shRNA seed sequences. cWords cluster plots showing the Z-scores of the 6mer, 7mer and 8mer nucleotide sequences within the top 10 enriched motifs in the 3’ UTR (left panels) or coding sequence (CDS, right panels) of genes downregulated by shTMEFF2-4 (**A**) and shTMEFF2-9 (**B**), shTMEFF2-9 according to RNA-seq analyses. X-axis contains rank ordered genes from the most downregulated to upregulated expression. Each mark represents an individual sequence, and red marks are sequences within the most enriched motifs. Triangles indicate seed sequences of known miRNAs (listed). Circles indicate sequences not currently known to correspond to human miRNA seeds.

**Figure 4 – figure supplement 3.**
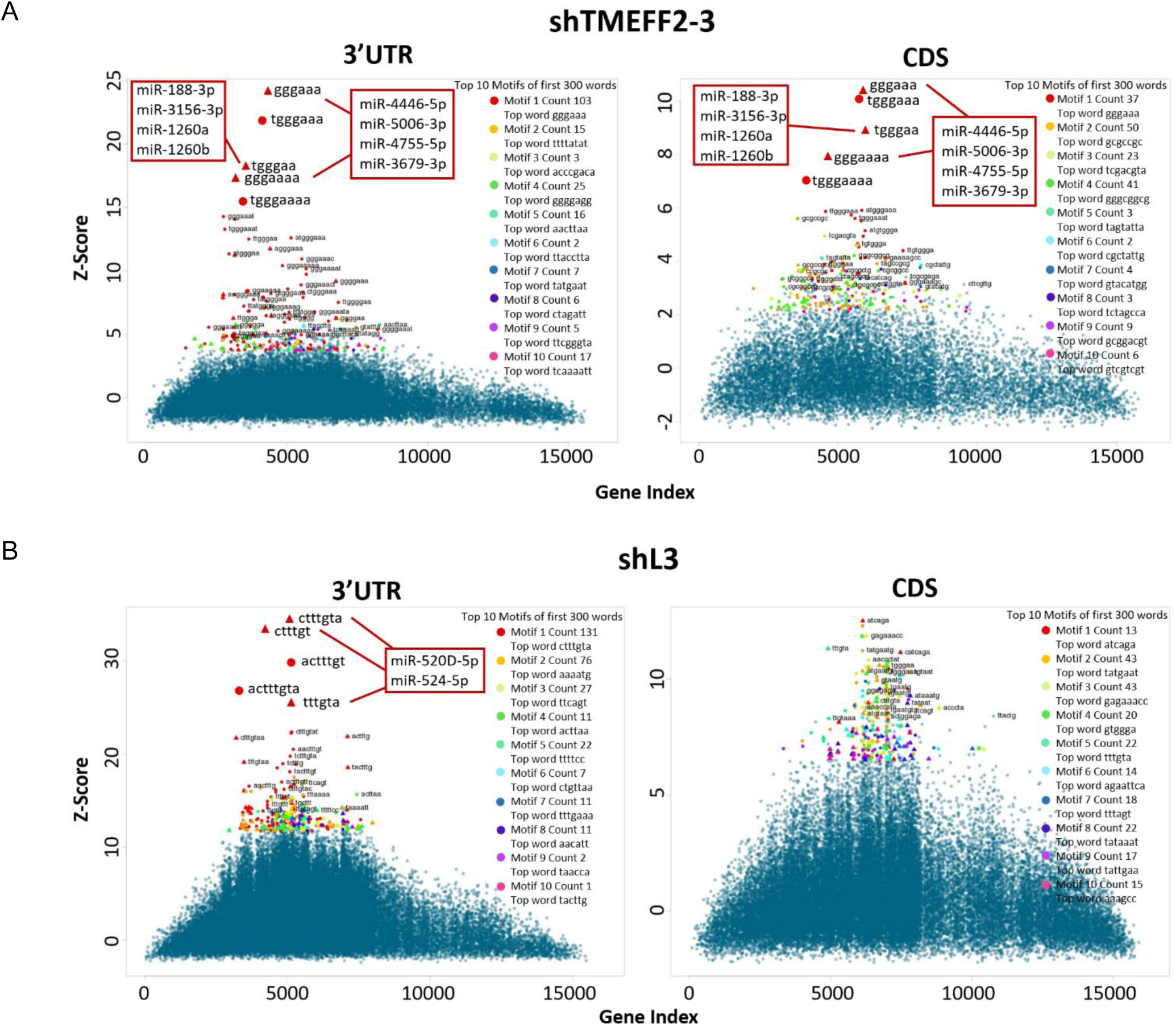
The top 3’ UTR motifs associated with shRNA-mediated gene downregulation are complementary to potential shRNA seed sequences. cWords cluster plots showing the Z-scores of the 6mer, 7mer and 8mer nucleotide sequences within the top 10 enriched motifs in the 3’ UTR (left panels) or coding sequence (CDS, right panels) of genes downregulated by shTMEFF2-3 (**A**) and shL3 (**B**) according to RNA-seq analyses. X-axis contains rank ordered genes from the most downregulated to upregulated expression. Each mark represents an individual sequence, and red marks are sequences within the most enriched motifs. Triangles indicate seed sequences of known miRNAs (listed). Circles indicate sequences not currently known to correspond to human miRNA seeds.

**Figure 4 – figure supplement 4.**
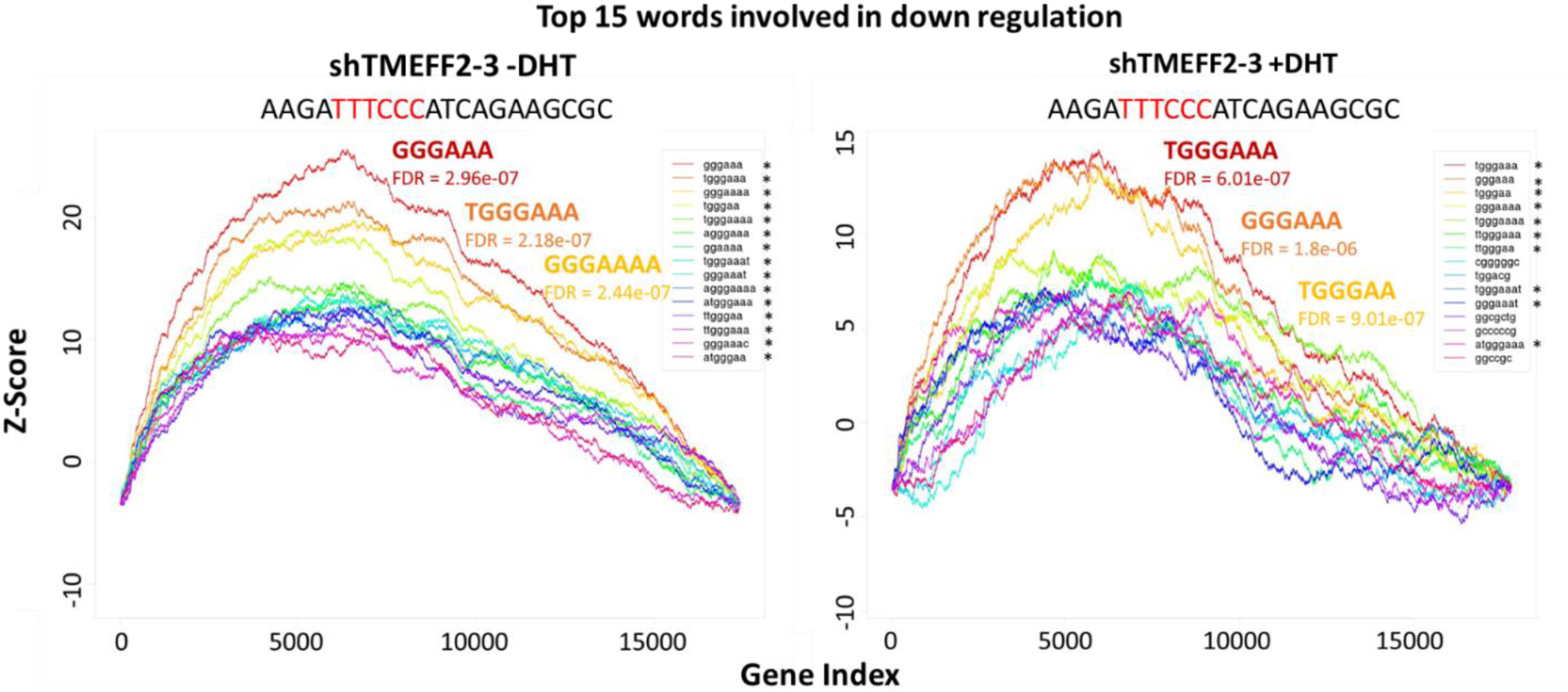
3’ UTR enriched sequences associated with shTMEFF2-3-mediated gene downregulation in cells grown in the presence and absence of DHT are complementary to potential shRNA seed sequences. (**A**) cWords enrichment plots showing the top 15 enriched 6mer, 7mer and 8mer in the 3’ UTR of genes downregulated by shTMEFF2-3 in the absence of DHT (left panel) and in the presence of DHT (right panel) according to RNA-seq analyses. Y-axis shows Z-score enrichment values. X-axis contains rank ordered genes from the most downregulated to upregulated expression. The top 3 enriched 3’ UTR sequences and FDR values are labeled (red: most enriched, orange: second most enriched, yellow: third most enriched). Unprocessed shTMEFF2-3 guide strand sequence is above each plot, and the potential 6mer seed sequence complementary to the most enriched 6mer 3’ UTR sequence associated with gene downregulation is in red. * indicates enriched sequences that are complementary to potential shTMEFF2-3 seed motifs.

**Figure 4 – figure supplement 5.**
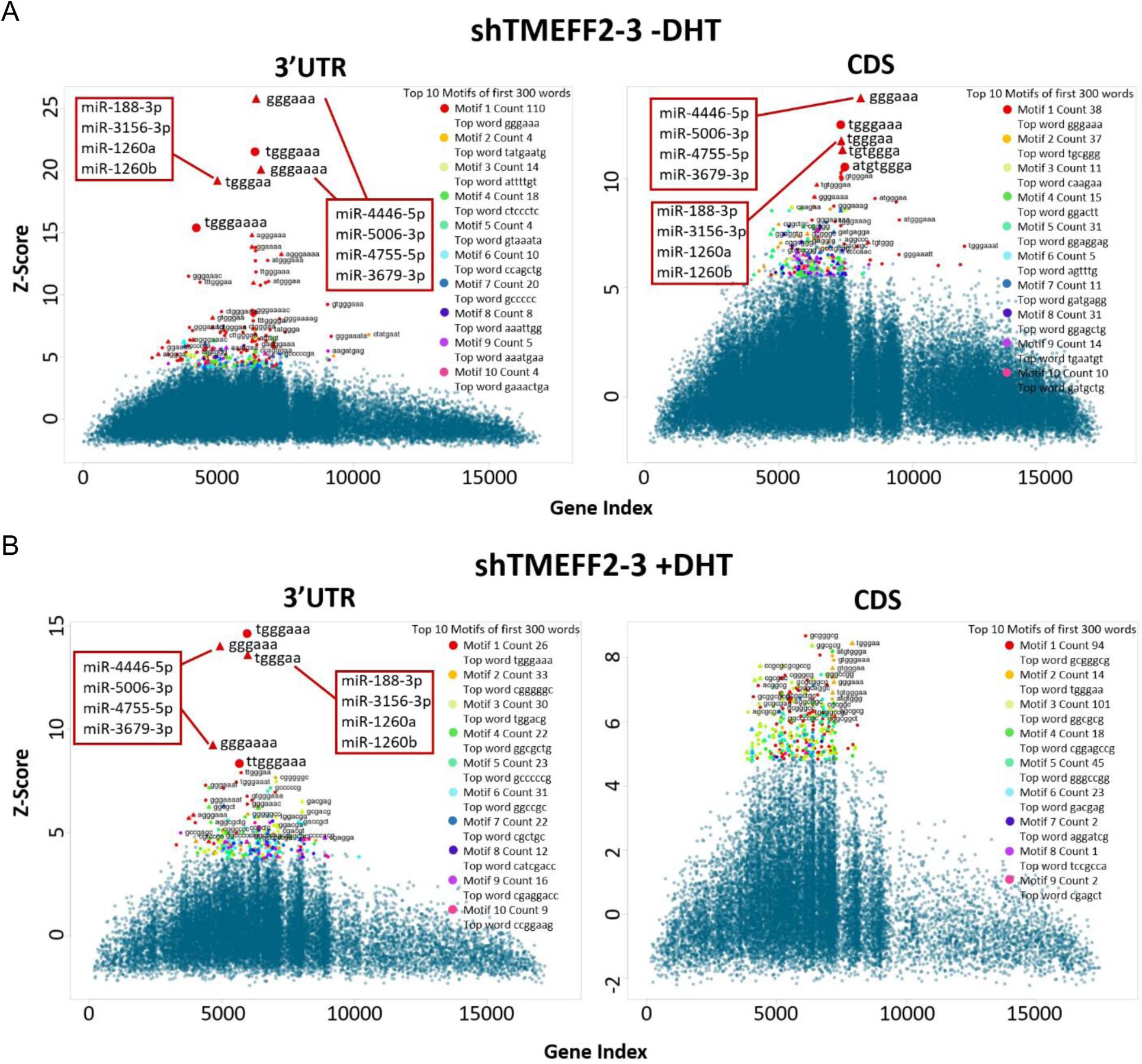
3’ UTR enriched sequences associated with shTMEFF2-3-mediated gene downregulation in cells grown in the presence and absence of DHT are complementary to potential shRNA seed sequences. cWords cluster plots showing the Z-scores of the 6mer, 7mer and 8mer nucleotide sequences within the top 10 enriched motifs in the 3’ UTR (left panels) or coding sequence (CDS, right panels) of genes downregulated by shTMEFF2-3 in the absence of DHT (**A**) and in the presence of DHT (**B**) according to RNA-seq analyses. X-axis contains rank ordered genes from the most downregulated to upregulated expression. Each mark represents an individual sequence (red marks are sequences within the most enriched motifs). Triangles indicate seed sequences of known miRNAs (listed). Circles indicate sequences not currently known to correspond to human miRNA seeds.

**Figure 4 – figure supplement 6.**
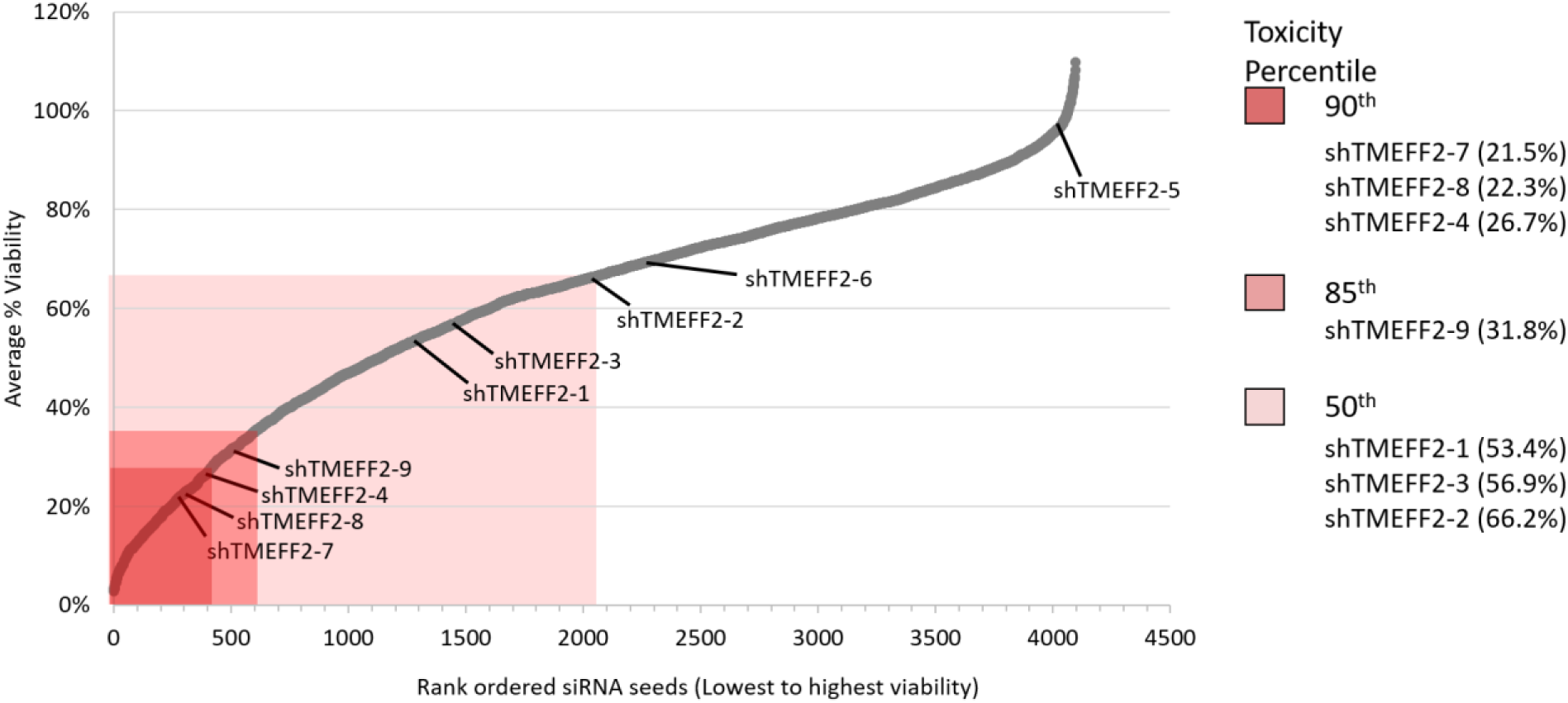
*In silico* analysis of siRNA seed viability screen reveals enrichment of potential TMEFF2 shRNA seeds within the most toxic seeds. Dot plot showing the average viability of HeyA8 and H460 cells transfected with siRNAs containing all possible (4096) 6mer seed sequences (data from Gao et al (Gao et al., 2018)). Seeds are rank ordered from effecting the lowest to highest viability and red rectangles mark percentile rankings of seed toxicity (90^th^, 85^th^, and 50^th^ percentile). Average percent viability of TMEFF2-targeted shRNA seeds, assuming consistent Dicer cuts during shRNA processing, are indicated only for those shRNA seeds within the 50^th^ percentile of toxicity.

**Figure 4 – figure supplement 7.**
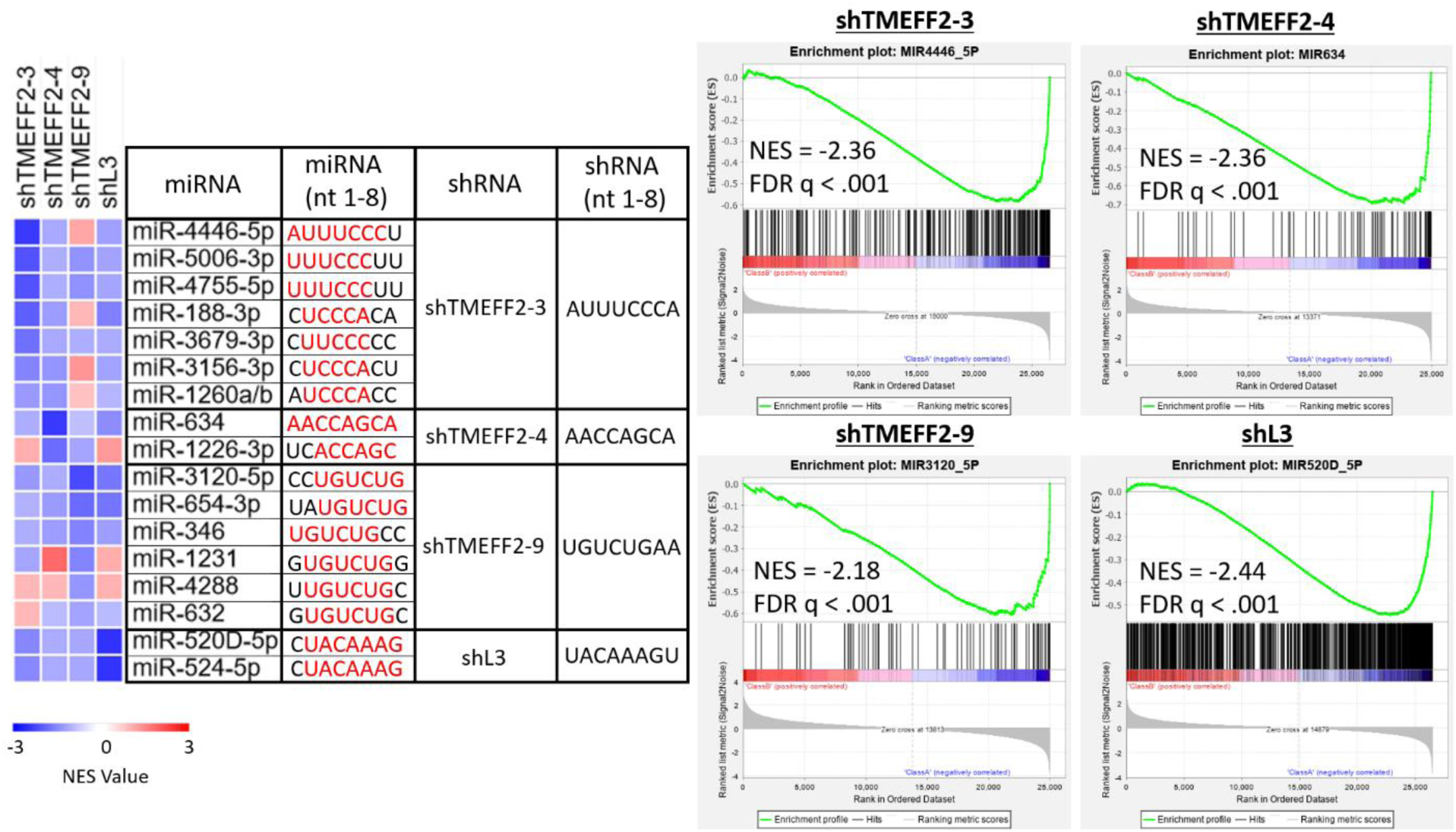
Gene sets of potential miRNA targets are downregulated by shRNAs with similar seed sequences. GSEAs of miRDB predicted target gene sets of miRNAs with seed motifs similar to shTMEFF2-3, -4, -9 or shL3 (as predicted by cWords analyses). Heatmap (left panel) shows the NES of each gene set. Positive NES indicates gene sets are upregulated, and negative NES indicates gene sets are downregulated by the designated shRNA. The first eight nucleotides of the guide strand of miRNAs and shRNAs are listed, and identical miRNA and shRNA sequences are in red. GSEA enrichment plots (right panel) showing the enrichment of the most significantly enriched miRDB gene set in rank ordered gene expression lists (upregulated to downregulated) for each shRNA. NES and FDR q-values are labeled on each plot.

**Figure 4 – figure supplement 8.**
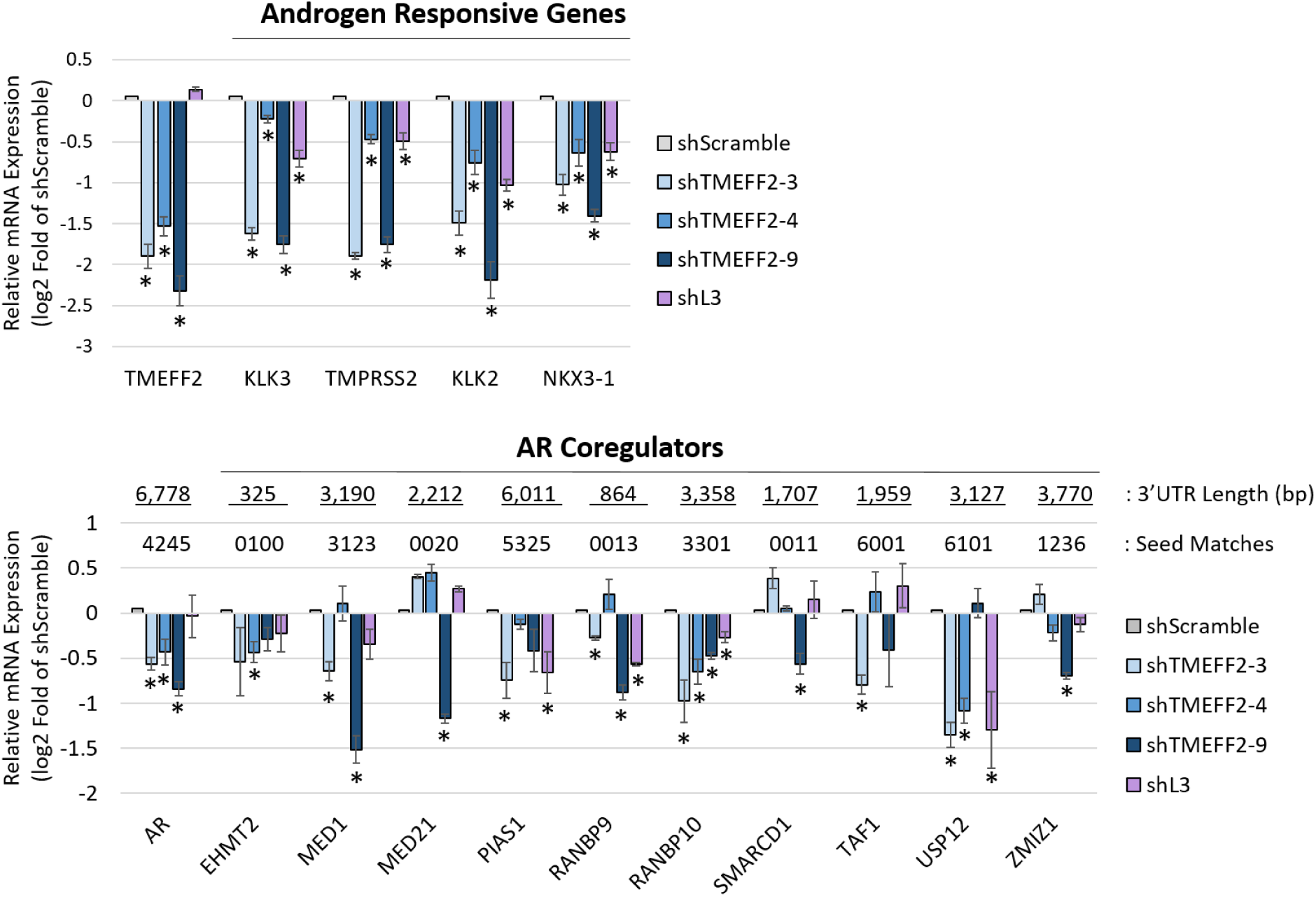
RT qPCR validation of androgen responsive and AR coregulatory gene downregulation by shRNAs. mRNA expression validation of androgen responsive (top panel) and AR coregulatory (bottom panel) gene expression. Genes selected were downregulated in LNCaP cells expressing the designated shRNAs based on RNA-seq analyses. Shown is the relative mRNA expression determined by RT qPCR (as log2 fold of shScramble). 3’ UTR length and number of 3’ UTR sequences complementary to 6mer seeds for each shRNA are labeled. RPL8, RPL38, PSMA1 and PPP2CA were housekeeping genes used for normalization. N=3, error bars ±SD, * p<.05 determined by t-test.

**Figure 4 – figure supplement 9.**
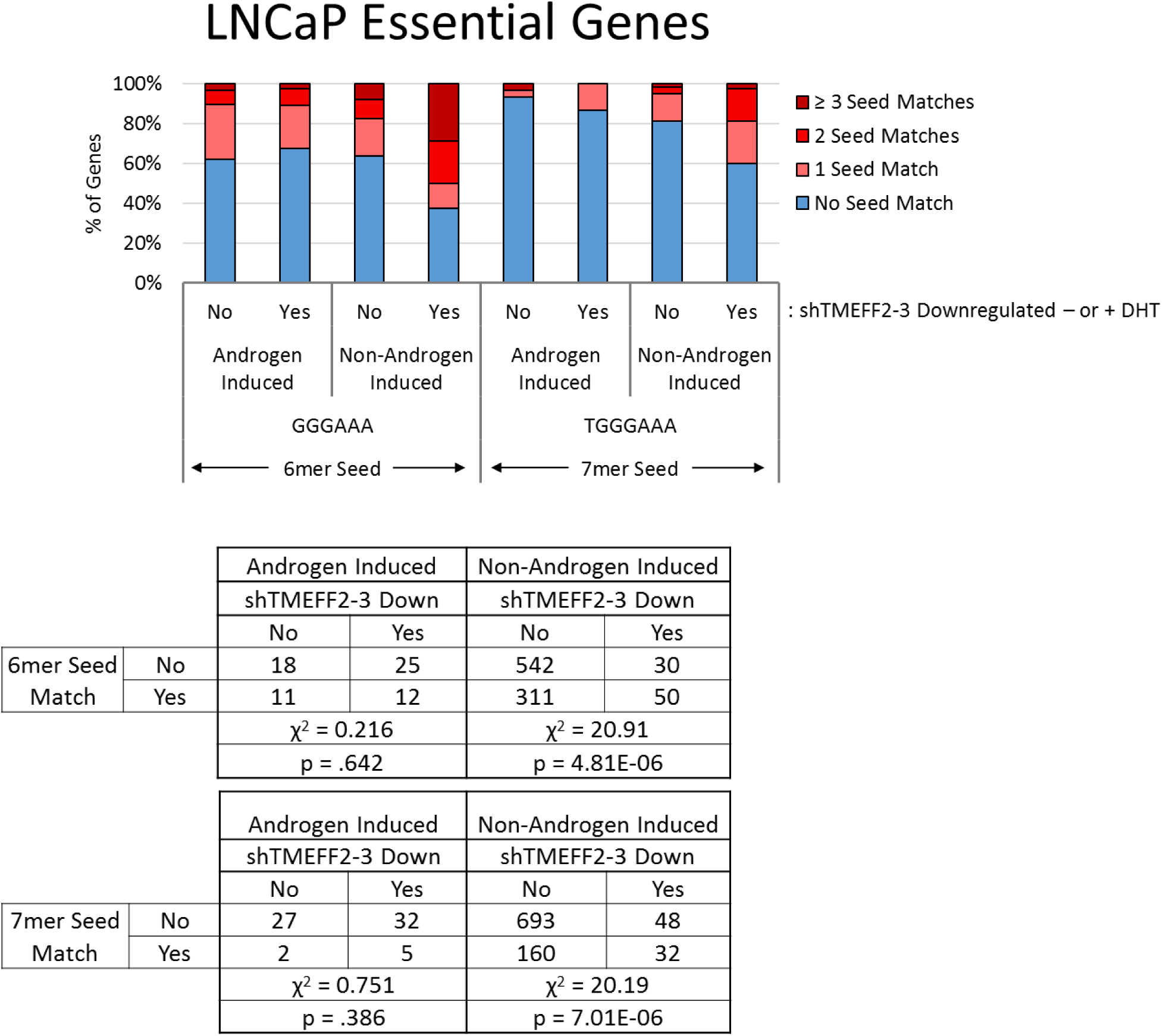
AR coregulatory and essential gene downregulation is associated with 3’ UTR complementary to sh-TMEFF2-3 seed sequence, and/or androgen signaling inhibition. LNCaP essential gene set (Fei et al., 2017) stratified by whether genes are induced by DHT (Androgen induced or Non-Androgen induced, based on data in shScramble cells), downregulation by shTMEFF2-3 (yes or no), and by the presence in their 3’-UTR of single, 2, or more, 6mer or 7mer sequences identified by cWords analyses. Only the most significantly associated with gene downregulation by shTMEFF2-3 (potential 3’ UTR seed matches) were used. Contingency tables are located below each stacked bar graph. P-values were calculated by chi square test of independence, and are labeled on the bottom of each contingency table.

**Figure 5 – figure supplement 1.**
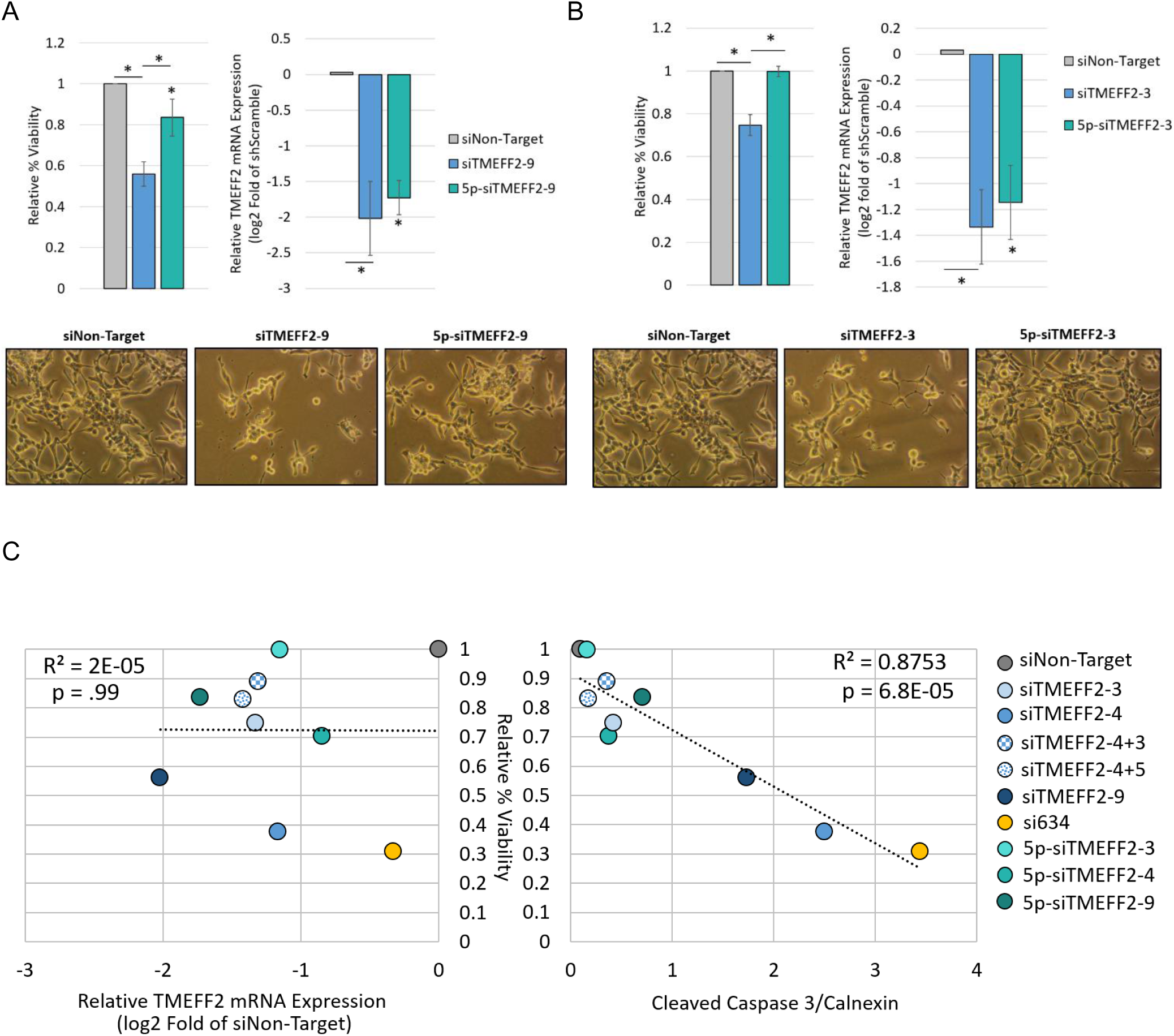
Reduction in PCa cancer cell viability is seed mediated. (**A/B**) Relative percent viability and TMEFF2 mRNA expression in LNCaP cells transfected with siTMEFF2-9 (**A**) or siTMEFF2-3 (**B**) and control siRNAs (siNon-Target; Negative seed control: 5p-siTMEFF2). 5p designates an ON-Target-Plus modification (Dharmacon) that blocks seed mediated gene downregulation. Cell viability was determined by trypan blue. Viability measurements, cell pictures and RNA extractions were done 72 hours after siRNA transfections. mRNA expression was determined by RT qPCR using RPL8, RPL38, PSMA1 and PPP2CA housekeeping genes for normalization. N=4, error bars ±SD, * p<.05 compared to siNon-target. Bars with * designate significant differences relative to siTMEFF2-9 or siTMEFF2-3. Significance was determined by t-test. (**C**) Correlation between relative percent viability and relative TMEFF2 mRNA expression (left) or normalized cleaved Caspase 3 (right) in LNCaP cells transfected with the designated siRNAs. See figure 5B for Caspase 3 western blot. Band intensity was quantified using Biorad Image Lab.

**Figure 5 – figure supplement 2.**
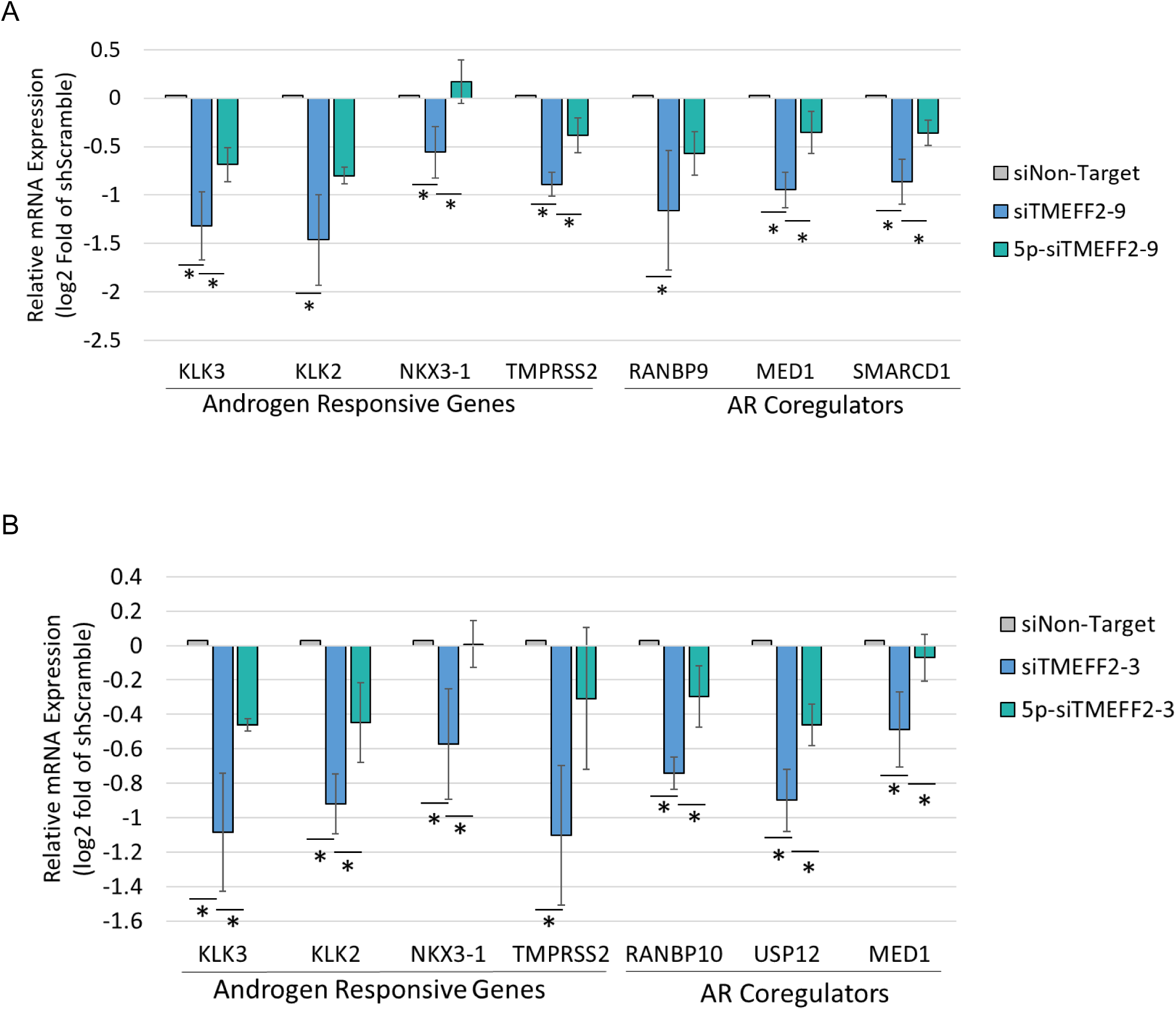
AR coregulatory gene downregulation are seed mediated. Androgen responsive, AR coregulator and AR mRNA expression in LNCaP cells transfected with siTMEFF2-9 (**A**) or siTMEFF2-3 (**B**) and control siRNAs (siNon-Target; Negative seed controls: 5p-siTMEFF2-3). 5p designates an ON-Target-Plus modification (Dharmacon) that blocks seed mediated gene downregulation. mRNA expression was determined by RT qPCR using RPL8, RPL38, PSMA1 and PPP2CA housekeeping genes for normalization. N=4, error bars ±SD, * p<.05 determined by t-test.

**Figure 5 – figure supplement 3.**
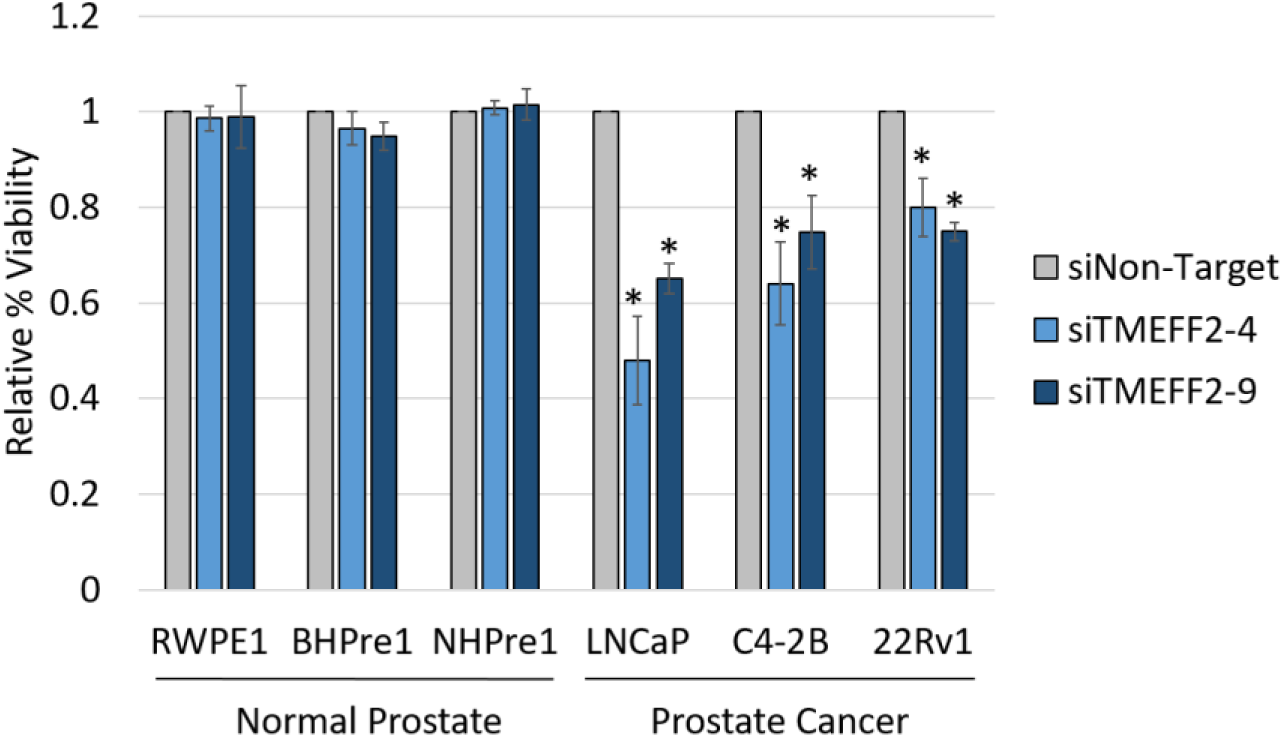
AN-DISE siRNAs do not reduce the viability of normal prostate epithelial cell lines. Relative percent viability of normal prostate (RWPE1, BHPre1, NHPre1) and PCa cell lines (LNCaP, C4-2B and 22Rv1) transfected with Cy5 labeled siNon-Target, siTMEFF2-4 or siTMEFF2-9 siRNAs. Viability was determined by trypan blue 72 hours after siRNA transfection. N=3, error bars ±SD, * p<.05 determined by t-test.

